# SNARE protein tomosyn regulates dense core vesicle composition but not exocytosis in mammalian neurons

**DOI:** 10.1101/2022.12.18.520925

**Authors:** Aygul Subkhangulova, Miguel A. Gonzalez-Lozano, Alexander J. A. Groffen, Jan R. T. van Weering, August B. Smit, Ruud F. Toonen, Matthijs Verhage

## Abstract

Tomosyn is a large, non-canonical SNARE protein proposed to act as a competitive inhibitor of SNARE complex formation in vesicle exocytosis. In the brain, tomosyn inhibits fusion of synaptic vesicles (SVs), whereas its role in the fusion of neuropeptide-containing dense core vesicles (DCVs) is unknown. Here, we addressed this question using a new mouse model allowing conditional deletion of tomosyn (*Stxbp5*) and its paralogue tomosyn-2 (*Stxbp5l*), and an assay that detects DCV exocytosis with single vesicle resolution in primary hippocampal neurons. Surprisingly, loss of both tomosyns did not affect DCV exocytosis but resulted in a strong reduction of intracellular levels of many DCV cargos, most prominently brain-derived neurotrophic factor (BDNF), granin VGF and prohormone convertase PCSK1. Reduced levels of DCV cargos were paralleled by decreased DCV size and impaired mRNA expression of the corresponding genes. We conclude that tomosyns regulate neuropeptide and neurotrophin secretion via control of DCV cargo production, and not at the step of cargo release. Our findings suggest a differential effect of tomosyn on the two main secretory pathways in mammalian neurons and argues against a conserved role of tomosyn as competitive inhibitor of SNARE complex formation.

## Introduction

Brain activity relies on the precisely regulated secretion of different chemical messengers, and most neurons co-release multiple neurotransmitters (Nusbaum *et al*, 2017). Classical small-molecule neurotransmitters (e.g., glutamate, γ-aminobutyric acid) are released from synaptic vesicles (SV), whereas neuropeptides and neurotrophic factors are packaged in and secreted from dense core vesicles (DCV). Exocytosis of both types of vesicles requires an identical basic machinery consisting of SNARE proteins: syntaxin-1 and SNAP25 at the plasma membrane, and VAMP2 on the vesicle. The SNARE domains of these proteins form a tight complex that brings the two membranes together and drives vesicle fusion. In addition to the basic SNARE machinery, many other proteins participate in the fusion process (reviewed in (Südhof, 2013; Rizo & Xu, 2015), of which some are differentially required for SV and DCV exocytosis (Persoon *et al*, 2019; Moro *et al*, 2021). The differences in molecular machinery may explain why SV and DCV exocytosis is triggered upon different stimulation paradigms and at different locations within neurons (reviewed in (Zupanc, 1996; Scalettar, 2006).

Tomosyn (STXBP5) is a large, evolutionary conserved SNARE protein discovered as an interactor of syntaxin-1 in rat brain (Fujita *et al*, 1998; Masuda *et al*, 1998). Tomosyn is expressed in a wide range of specialized secretory cell types, such as neurons, platelets, and neuroendocrine cells, suggesting a fundamental role of tomosyn in regulated secretion. Mammals express a second tomosyn gene, *STXBP5L*, which encodes tomosyn-2 with a nearly 80% amino acid similarity (Groffen *et al*, 2005). The two genes have a partially overlapping expression pattern in mouse brain suggesting functional redundancy (Groffen *et al*, 2005; Barak *et al*, 2010). Mutations in both genes were found in patients with neurodevelopmental disorders, suggesting a role of tomosyns in human brain function (Matsunami *et al*, 2013; Cukier *et al*, 2014; De Rubeis *et al*, 2014; Kumar *et al*, 2015).

Tomosyn is comprised of a C-terminal SNARE domain and N-terminal WD40 repeats folded into two seven-bladed β-propellers. The SNARE-domain of tomosyn is homologous to the SNARE domain of VAMP2 and mediates binding to syntaxin-1 and SNAP25. The resulting complex is remarkably similar in structure and stability to the fusogenic complex of syntaxin-1/SNAP25 with VAMP2 (Pobbati *et al*, 2004). However, in contrast to VAMP2 and most other SNARE proteins, tomosyn does not contain a transmembrane anchor and therefore cannot drive membrane fusion. This feature led to a hypothesis that tomosyn functions as an inhibitor of fusion by competing with VAMP2 for binding to syntaxin-1/SNAP25 (Hatsuzawa *et al*, 2003). In support of this hypothesis, overexpression of tomosyn in neuroendocrine PC12 cells and adrenal chromaffin cells inhibits fusion of secretory vesicles (Fujita *et al*, 1998; Hatsuzawa *et al*, 2003; Yizhar *et al*, 2004), and, conversely, loss of tomosyn in nematode, fruit fly and mouse brain increases SV release probability and enhances synaptic transmission (Gracheva *et al*, 2006; McEwen *et al*, 2006; Chen *et al*, 2011; Sauvola *et al*, 2021; Sakisaka *et al*, 2008; Ben-Simon *et al*, 2015). In addition, loss of tomosyn in *C. elegans* augments neuropeptide secretion, as evidenced by decreased DCV abundance at synapses and increased accumulation of a DCV cargo in scavenger cells (Gracheva *et al*, 2007; Ch’ng *et al*, 2008). Together, these studies suggest an evolutionary conserved role of tomosyn as an inhibitor of secretory vesicle exocytosis.

However, several lines of evidence argue against this conclusion. First, rat sympathetic neurons show a decrease in neurotransmitter release upon depletion of tomosyn (Baba *et al*, 2005). Second, platelets from *Stxbp5*-null mice show severely reduced exocytosis of all types of secretory granules, which results in excessive bleeding and impaired thrombus formation *in vivo* (Zhu *et al*, 2014; Ye *et al*, 2014). Finally, loss of SRO7 and SRO77, two yeast orthologs of mammalian tomosyns, causes a strong defect in exocytosis of post-Golgi vesicles (Lehman *et al*, 1999). Collectively, these studies suggest a positive role of tomosyn and its yeast orthologs in exocytosis. Thus, despite the clear importance of tomosyn in regulated secretion, its role in this process remains controversial.

In the brain, the role of mammalian tomosyn has been studied with a focus on SV exocytosis, whereas its effect on DCV exocytosis is unknown. In this study, we addressed this question using a mouse model with the inducible loss of all tomosyn and tomosyn-2 isoforms and show that tomosyns do not affect neuronal DCV exocytosis but, paradoxically, regulate DCV composition and intracellular levels of brain-derived neurotrophic factor (BDNF) and many neuropeptides. Thus, tomosyns differentially modulate two regulated secretory pathways in neurons (secretion from SVs and DCVs) and have an additional role in the regulation of intracellular DCV cargo levels.

## Results

To explore the role of tomosyn in neuronal secretion, we generated a new mouse model to allow conditional deletion of tomosyn (*Stxbp5*) and tomosyn-2 (*Stxbp5l*) (Figure 1 supplement 1). Primary hippocampal neurons isolated from these mice showed a complete loss of tomosyns expression when transduced with a lentivirus encoding *Cre*-recombinase, as shown by Western blot (WB) and immunocytochemistry (ICC). As wildtype (WT) control, we used neurons isolated from the same litter but expressing a defective *Cre*-recombinase that lacks the DNA-binding domain (ΔCre). Double knockout (DKO) neurons developed normally in culture and did not show any defects in neurite outgrowth and synapse formation as evidenced by normal dendrite length and density of synaptophysin puncta (Figure 1 supplement 1 D-G).

To examine DCV fusion, we used a modification of a validated reporter consisting of neuropeptide Y (NPY) fused to super-ecliptic pHluorin (Figure 1A, Figure 1 supplement 2) (Persoon *et al*, 2018; Nassal *et al*, 2022). When expressed in primary neurons, this reporter localized almost exclusively to DCVs, as indicated by its co-localization with endogenous DCV markers IA-2 and chromogranins (CHGA, CHGB) (Figure 1 supplement 2). Fusion of labeled DCVs with the plasma membrane was stimulated by repetitive bursts of action potentials (50Hz) delivered to single neurons grown on glial micro-islands. As shown previously (Persoon *et al*, 2018; van Westen *et al*, 2021), this stimulation resulted in a steep accumulation of fusion events that plateaued shortly before the end of the stimulation (Figure 1B). Surprisingly, the total number of fusion events was similar between WT and DKO neurons, with a tendency (*p*=0.09) towards a decrease in DKO neurons (Figure 1C). However, the total intracellular pool of the DCV reporter, visualized by a short application of 50 mM NH_4_Cl to neurons to dequench intravesicular NPY-pHluorin, was decreased by 28% in DKO neurons (Figure 1D). The released fraction, i.e. the number of fusion events normalized to the intracellular pool of vesicles, was nearly identical between WT and DKO neurons (Figure 1E). Also, the fluorescence peak of individual fusion events, an estimate of NPY loading per vesicle, was not affected by the loss of tomosyns (Figure 1F). Thus, tomosyns are dispensable for the fusion of neuronal DCVs but regulate intracellular levels of the DCV reporter.

**Figure 1.**
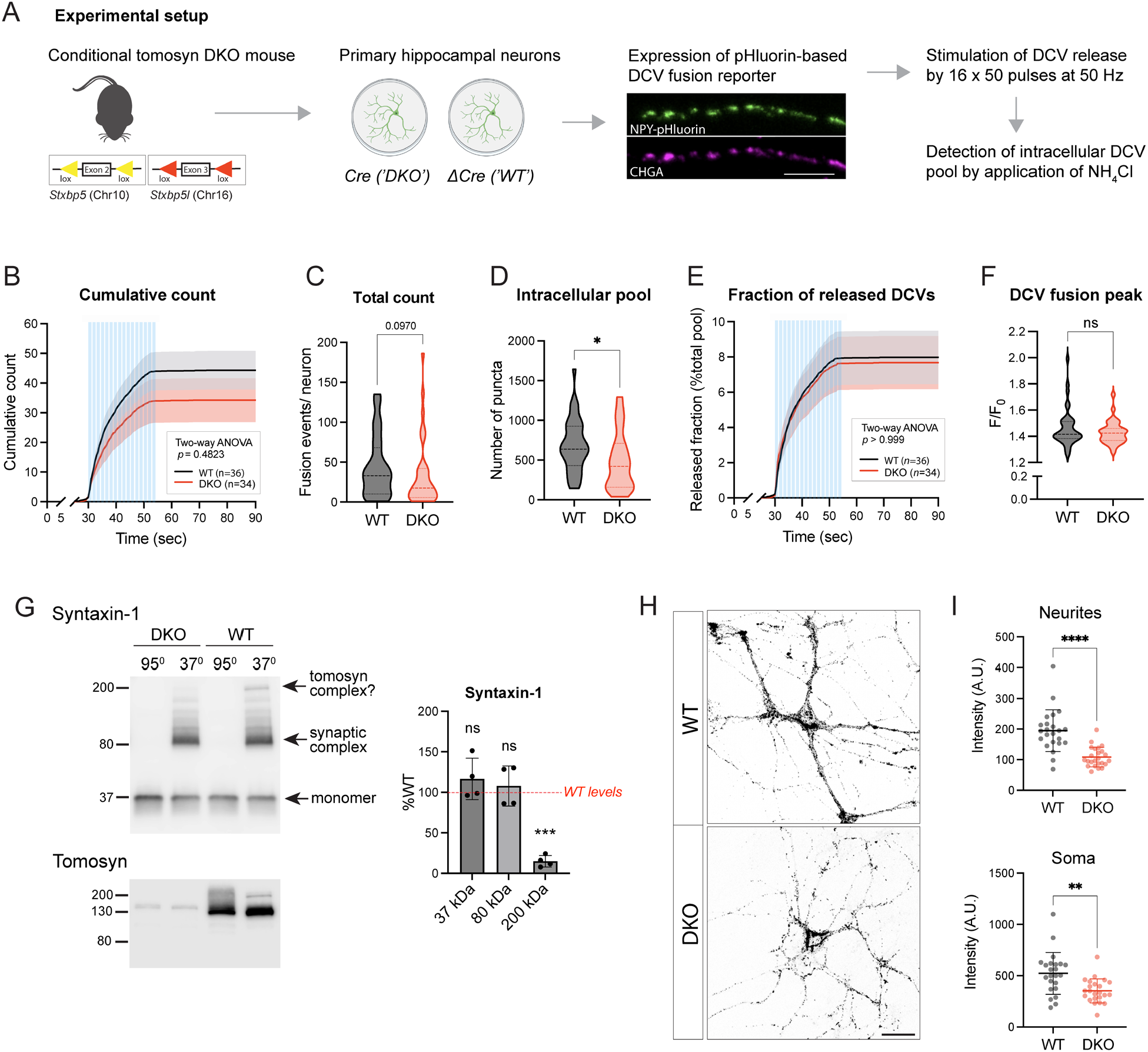
Loss of tomosyns does not affect fusion of DCVs in mouse hippocampal neurons. (A) Experimental strategy for measuring DCV fusion in single isolated neurons. Results of the experiment are shown in (B-E). (B) Cumulative count of fusion events detected per single neuron. Blue lines indicate time of electric stimulation (16 bursts of 50 pulses at 50 Hz each). Data are shown as mean ± SEM and were analyzed by two-way ANOVA. n=34-36 neurons/ genotype from three culture preparations. (C) Total count of fusion events detected per single neuron. Data are shown as violin plot with median (dashed line) and quartiles (dotted lines) and analyzed using Mann-Whitney test. (D) Intracellular pool of pHluorin-labeled DCVs determined by application of 50 mM NH_4_Cl after the stimulation. Data are shown as violin plot with median (dashed line) and quartiles (dotted lines) and analyzed using a two-tailed unpaired *t*-test. \**p* < 0.05. (E) Cumulative count of fusion events normalized to the intracellular pool of DCVs (release fraction). Data are shown as mean ± SEM and were analyzed by two-way ANOVA. (F) Peak in fluorescence intensity that results from a single fusion event (averaged per neuron). Data are shown as violin plot with median (dashed line) and quartiles (dotted lines) and analyzed using Mann-Whitney test. ns: not significant. (G) Levels of syntaxin-1 containing 80 kDa SNARE complexes are unchanged in DKO neurons. Boiled and non-boiled lysates were subjected to SDS-PAGE and membranes were probed with an antibody against syntaxin-1 (STX1). Temperature-sensitive complexes were detected at 80 kDa (synaptic SNARE complex) and 200 kDa (possibly complex of syntaxin-1 and SNAP25 with tomosyn). Smear at >80 kDa represents complex multimers. Band intensities in DKO were normalized to WT, plotted as mean ± SD and analyzed using a one sample *t*-test. Dots represent independent culture preparations (n=4). ns: not significant; \*\*\**p* < 0.001. (H) DCV reporter levels are reduced in DKO neurons silenced with the sodium channel blocker tetrodotoxin (TTX). Representative images of NPY-pHluorin in fixed WT and DKO neurons are shown. Neurons were grown in network (mass) cultures, treated with 1 µM TTX for 48 hours, and fixed on DIV14 before imaging. Scale bar 20 µm. (I) Mean intensity of the DCV reporter in neurites and somas of silenced WT and DKO neurons exemplified in (H). Data are shown as mean ± SD and were analyzed using a two-tailed unpaired *t*-test with Welch’s correction. n=23-24 neurons/ genotype. Dots represent individual neurons. \*\**p* < 0.01, \*\*\*\**p* < 0.0001.

Tomosyn was proposed to act as an inhibitor of vesicle fusion by competing with VAMP2 for the binding to syntaxin-1 and SNAP25. Hence, loss of tomosyns may result in the increased formation of ternary synaptic SNARE complexes consisting of syntaxin-1, SNAP25 and VAMP2, as was shown for tomosyn-deficient fly brains (Sauvola *et al*, 2021). We examined levels of the assembled complexes in DKO neurons by SDS-PAGE on non-boiled neuronal lysates, based on the fact that these complexes are resistant to SDS but are disassembled at temperatures higher than 60°C (Hayashi *et al*, 1994; Otto *et al*, 1997) (Figure 1G). As expected, a band of approximately 80 kDa that corresponds to the ternary complexes was readily detected with a syntaxin-1 antibody in non-boiled lysates and absent in lysates heated at 95°C. No difference in the intensity of the 80 kDa band was observed between the genotypes. These data suggest that tomosyns do not interfere with the formation of syntaxin-1 containing synaptic SNARE complexes in cultured neurons. Interestingly, in addition to the 80 kDa band, non-boiled WT lysates showed a band of approximately 200 kDa, which was absent in DKO lysates and, therefore, likely represents a complex of tomosyn with syntaxin-1 and SNAP25, suggesting that this complex is resistant to SDS *in vivo*. This 200 kDa complex was also detected in WT neurons with an antibody against tomosyn (Figure 1G).

Despite normal DCV exocytosis, intracellular levels of the DCV reporter were decreased in DKO neurons (Figure 1D). This decrease was also observed in tetrodotoxin (TTX)-silenced network cultures of DKO neurons that were not subjected to electric stimulation (Figure 1H-I). Activity of the synapsin promoter used to drive expression of the DCV reporter was not significantly altered in DKO neurons, as shown by normal expression of enhanced green fluorescent protein (EGFP) from the same promoter (Figure 1 supplement 3). These data suggest that the decreased levels of the DCV reporter are not caused by decreased transcriptional activity of the construct driving the reporter expression.

We next examined if endogenous DCV cargos are also affected by the loss of tomosyns. Levels of brain-derived neurotrophic factor (BDNF), one of the most common DCV cargos in excitatory hippocampal neurons (Dieni *et al*, 2012), were decreased by 47% in DKO neurons as shown by ICC (Figure 2A-B). Levels of IA-2 (PTPRN), a transmembrane DCV-resident protein commonly used as a DCV marker, were also lower, by 25%, in DKO neurons as indicated by ICC, and by approximately 50% as evidenced by WB analysis, which allows to distinguish a proteolytically processed (mature) form of IA-2 that runs at approximately 65 kDa (Hermel *et al*, 1999) (Figure 2C-E). The cellular distribution of the two DCV markers was similar between the genotypes, with a comparable reduction in somas and neurites of DKO neurons and no re-distribution between the two compartments as detected by ICC. Markers of lysosomes/late endosomes, LAMP1 and LIMP2, were not affected by the loss of tomosyns, as shown by ICC and WB (Figure 2 supplement 1). These data suggest that tomosyns regulate the number and/or composition of neuronal DCVs but not of other post-Golgi vesicles.

**Figure 2.**
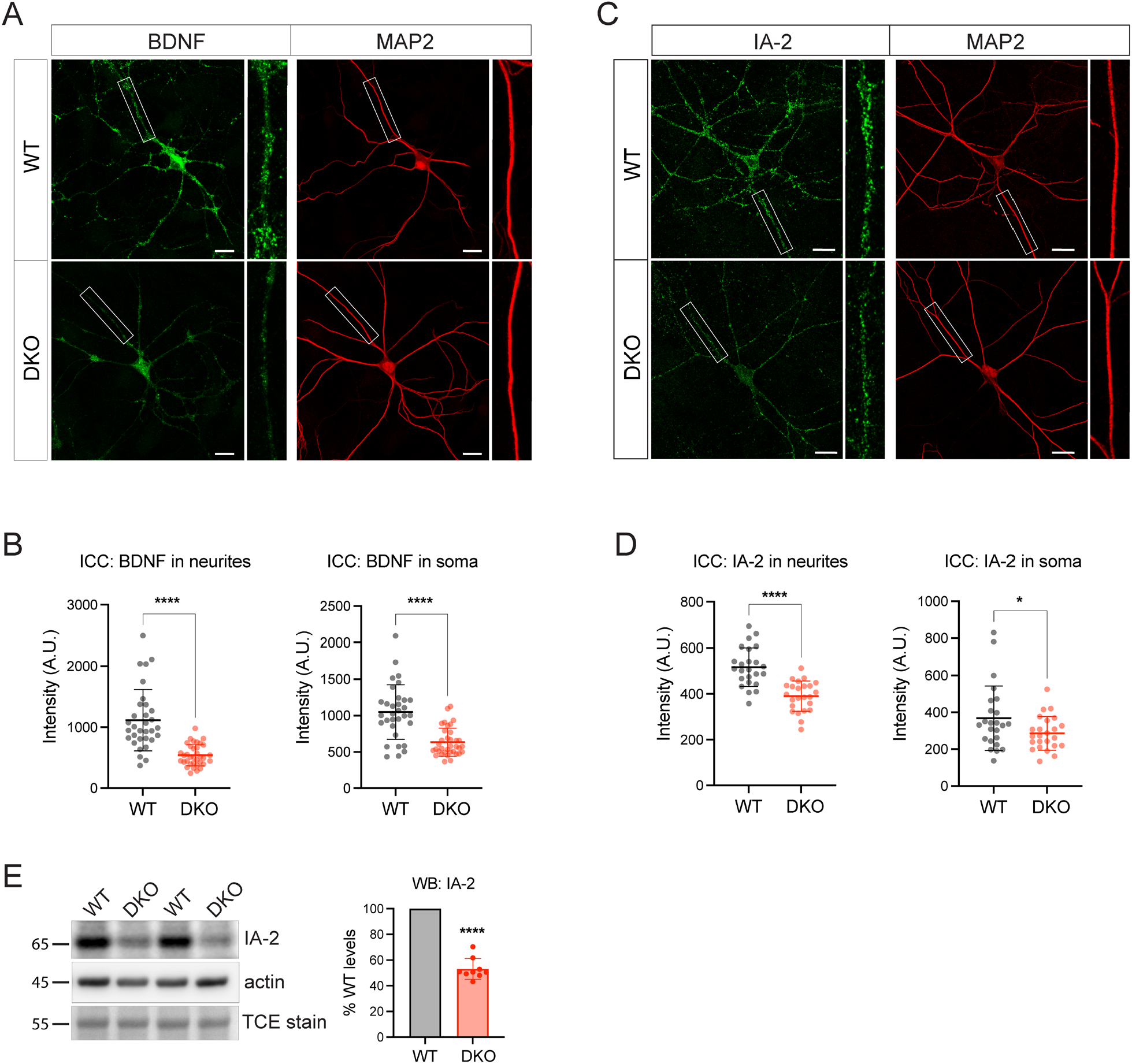
Tomosyns regulate number and/ or composition of neuronal DCVs. (A) BDNF levels are decreased in DKO neurons. Representative images of BDNF immunostaining in single DIV21 neurons grown on glia microislands. White boxes indicate a zoomed view of the neurites with BDNF puncta. Scale bar 20 µm. (B) Quantification of the mean BDNF intensity in WT and DKO neurons from confocal microscopy images exemplified in (A). Data are shown as mean ± SD and were analyzed using a two-tailed unpaired *t*-test. n=32-35 neurons/ genotype from three culture preparations. \*\*\*\**p* < 0.0001. (C) Levels of a DCV-resident protein IA-2 (PTPRN) are decreased in DKO neurons. Representative images of the IA-2 immunostaining in DIV14 neurons. White boxes indicate a zoomed view on the neurites with IA-2 puncta. Scale bar 20 µm. (D) Quantification of the mean IA-2 intensity in WT and DKO neurons from confocal microscopy images exemplified in (D). Data are shown as mean ± SD and were analyzed using a two-tailed unpaired *t*-test. n=24 neurons/ genotype. \**p* < 0.05; \*\*\*\**p* < 0.0001. (E) Levels of IA-2 as detected by WB are decreased in DKO neurons. Proteolytically processed (mature) form of IA-2 was detected at 65 kDa. Equal loading was verified by the detection of total protein stained with trichloroethanol (TCE) and by immunodetection of actin. Amount of mature IA-2 in DKO neurons was normalized to WT levels in the corresponding culture. DKO data are presented as mean ± SD and were analyzed using one sample *t*-test. n=9 samples/ genotype from three culture preparations. \*\*\*\**p* < 0.0001.

To analyze proteins affected by the loss of tomosyns in an unbiased manner, we compared the proteome of cultured WT and DKO neurons by mass spectrometry. We detected approximately 8500 proteins, of which around 3% were differentially expressed (log_2_ fold change threshold 0.3) (Figure 3A). Gene ontology (GO) analysis of the proteins decreased in DKO neurons indicated a rather selective downregulation of proteins related to the cytoskeleton and secretory granules (Figure 3B). Most of the up-regulated proteins were mitochondrial proteins. BDNF, along with the gene products of the perturbed genes (STXBP5 and STXBP5L), was detected only in WT but not in DKO neurons (Figure 3C). Other common DCV cargos were decreased to a various extent in DKO, among them: granins VGF and SCG2, cargo processing enzymes PCSK1 and CPE, and transmembrane DCV proteins IA-2 (PTPRN) and PAM (Figure 3D). In contrast to the DCV-resident proteins, SV proteins did not follow any general trend in DKO neurons: while some proteins (VGLUT2, GAD67) were strongly increased in DKO, others (e.g. SYT12 and SV2B) were reduced to a similar extent and many were not significantly altered. In conclusion, interfering with expression of both tomosyns leads to a reduction in the cellular levels of cytoskeletal and DCV cargo proteins and an increase in mitochondrial proteins.

**Figure 3.**
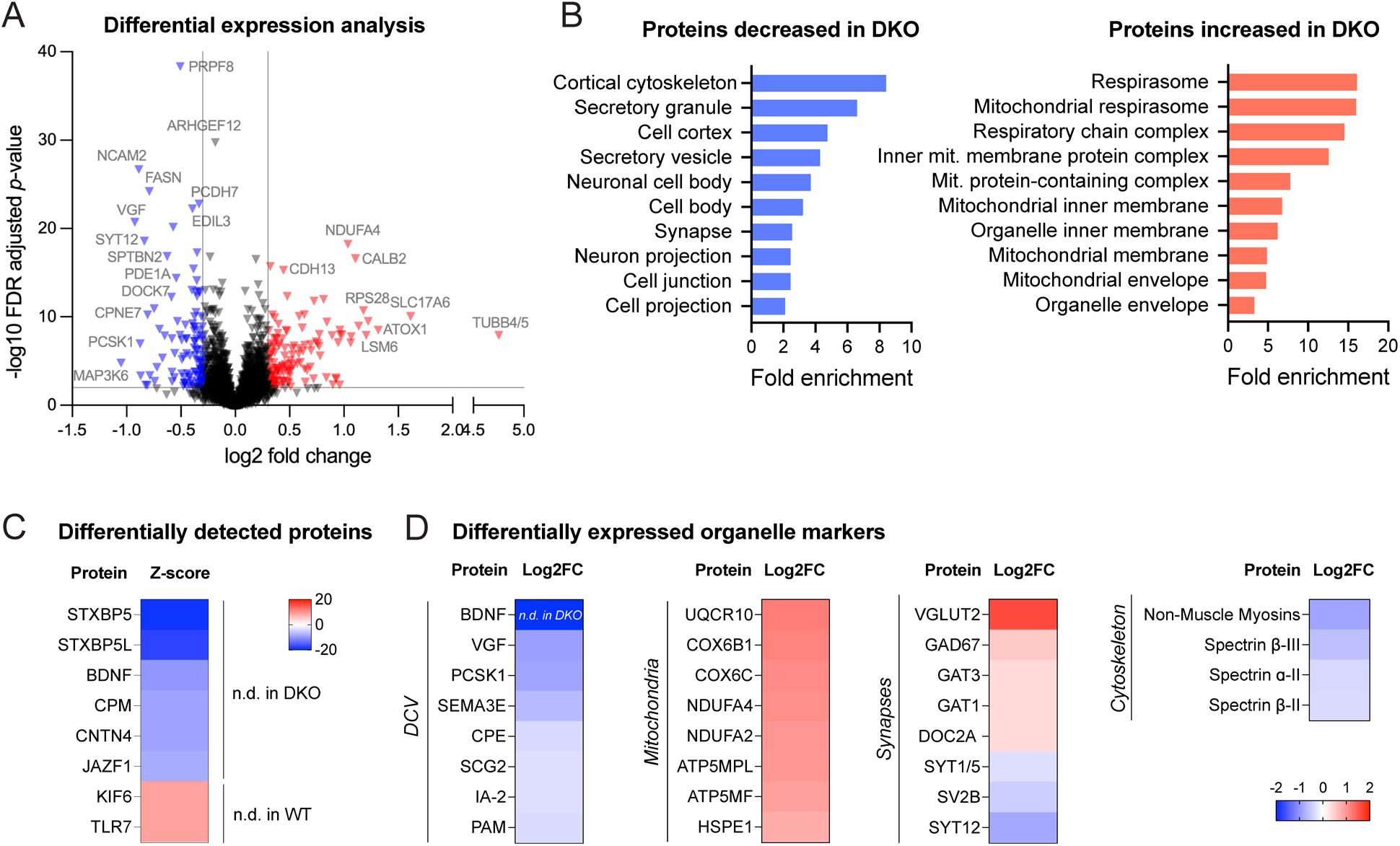
Proteomic analysis shows downregulation of DCV and cytoskeletal proteins in DKO neurons. (A) Volcano plot showing results of the differential expression analysis of WT and DKO proteome. Grey lines indicate chosen thresholds for log2 fold change (0.3) and false discovery (FDR) adjusted *p*-value (0.01). Proteins decreased or increased in DKO are shown in blue and red, respectively. (B) Enriched gene ontology (GO cellular component) terms of the proteins significantly affected in DKO neurons as analyzed by ShinyGO v0.75 (http://bioinformatics.sdstate.edu/go/) with the FDR-adjusted *p*-value cutoff ≤ 0.05. (C) List of proteins reliably detected in one of the genotypes only, and therefore not included in the differential expression analysis shown in (A). Proteins are sorted by the z-score, which reflects the total number of peptides detected per genotype. (D) Examples of proteins that are significantly affected in DKO neurons grouped by subcellular compartment. Heat maps show the degree of down-or upregulation in the DKO.

Next, we examined ultrastructural morphology of DCVs in DKO and WT neurons by electron microscopy (Figure 4). We found DCVs with normal dense cores sparsely distributed among neurites in neurons of both genotypes. Most presynaptic sections did not contain any DCVs, in line with previous reports (Sorra *et al*, 2006; Persoon *et al*, 2018). Percent of synaptic sections containing one or several DCVs was decreased in DKO by 30% (12% of all synaptic sections in DKO versus 17% in WT), indicating lowered DCV abundancy in DKO. In addition, DCVs in DKO neurons were slightly reduced in diameter (average 65.01 ± 2.08 nm in DKO versus 71.52 ± 1.62 nm in WT), with a high proportion of DKO DCVs being only 40-50 nm in diameter (16% of all DCVs in DKO versus 4% in WT) (Figure 4A-B). Such a reduction in diameter translates to an approximately 25% decrease in volume. In contrast, the diameter of SV was not affected by the loss of tomosyns (Figure 4D-E). Other parameters of synaptic ultrastructure, such as SV number per synapse, length of the active zone, and length of postsynaptic density, were also unchanged in DKO neurons (Figure 4D, 4F-H) and DKO somata showed no obvious abnormalities in organelle morphology (Figure 4D). Hence, loss of tomosyns leads to a specific reduction in DCV size and abundancy, without affecting ultrastructure and number of SVs.

**Figure 4.**
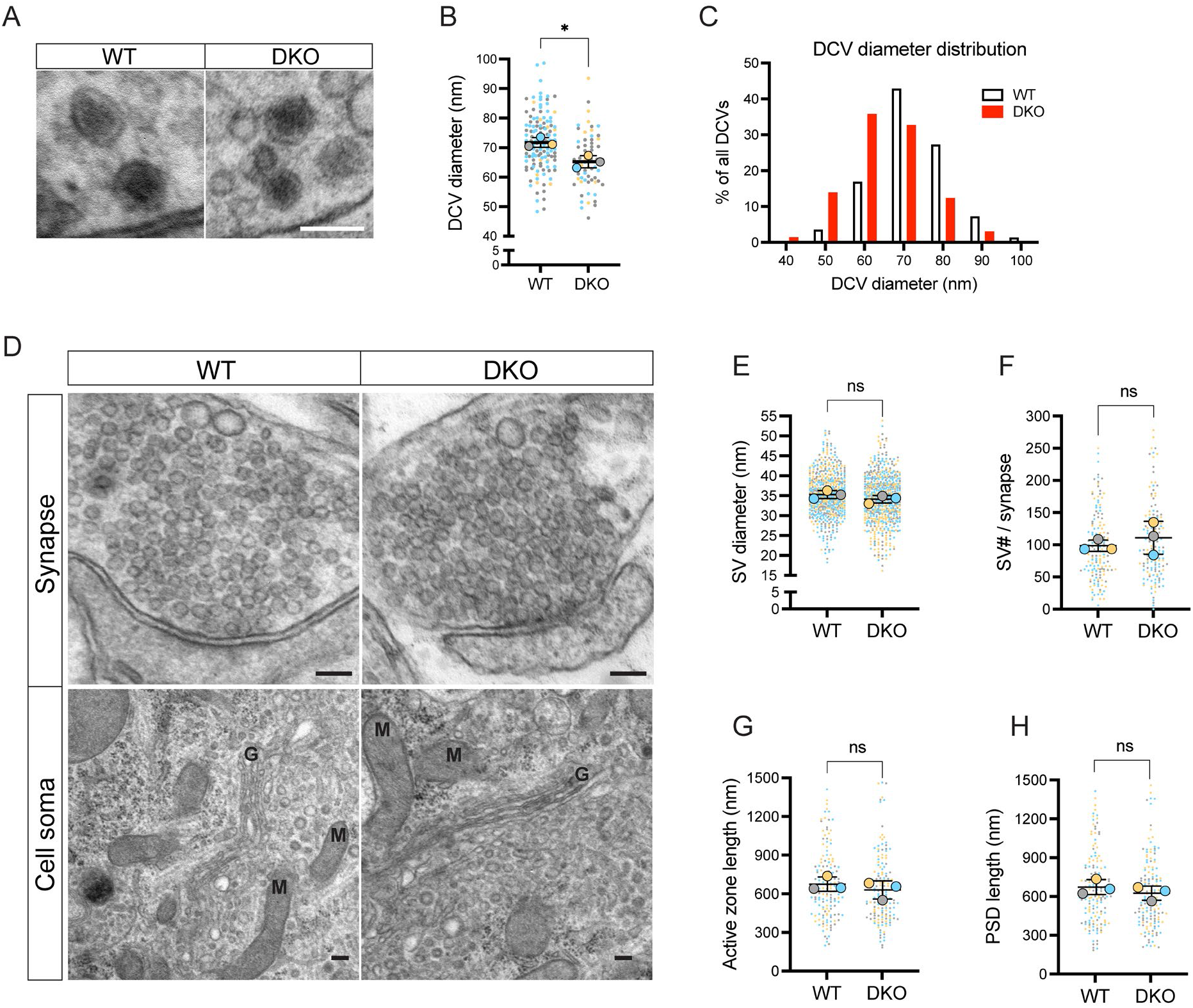
Loss of tomosyns results in a decrease of DCV but not of SV size. (A) Example EM images show normal appearance of DCVs with a typical dense core in DKO neurons. Scale bar 100 nm. (B) DCV diameter is reduced in DKO neurons, as quantified from EM images exemplified in (A). Data are shown as a SuperPlot, where averages from three independent cultures (biological replicates) are shown as large circles and single observations (DCV profiles) as dots. Data from each culture are shown in different color. Horizontal bars represent means of the averages from the three cultures, which were compared using a two-tailed unpaired *t*-test. Error bars are SD. Number of DCV profiles analyzed: 135 (WT) and 64 (DKO). \**p* < 0.05. (C) Frequency distribution of DCVs by diameter is skewed to the left in DKO neurons, indicating a higher proportion of smaller DCVs in DKO neurons than in WT. (D) Example EM images show normal ultrastructure of synapses and other cellular organelles in DKO neurons. Golgi cisternae and selected mitochondria are labeled with ‘G’ and ‘M’, respectively. Scale bar 100 nm. (E-H) Quantification of SV diameter, SV number per synapse, length of the active zone, and length of postsynaptic density (PSD) from EM images exemplified in (D). Data are shown as SuperPlots and analyzed as described in (B). Number of individual observations: 617-621 SV profiles (E) and 148-152 synapses (F-H). ns: not significant.

The fact that abundancy, composition, and size of neuronal DCVs is affected in DKO neurons suggests that tomosyns may regulate DCV biogenesis. DCVs are formed at the trans-Golgi network (TGN) in a sequence of membrane fusion/ fission events ensuring correct packaging and processing of DCV cargos. Interestingly, tomosyn is enriched at the TGN (Figure 5 supplement 1) and the TGN area is reduced by 15% in DKO neurons, as shown by immunostaining of the TGN marker TMEM87A (Figure 5A-B). We therefore tested whether tomosyn regulates packaging and export of a neuropeptide cargo from the TGN using a modification of the Retention Using Selective Hooks (RUSH) method (Boncompain *et al*, 2012). In this method, streptavidin-binding peptide (SBP)-tagged secretory cargo is retained in the endoplasmic reticulum (ER) until addition of biotin due to the expression of the ER-localized streptavidin. Biotin disrupts SBP/streptavidin interaction and induces a synchronized transport of the cargo from the ER along the secretory pathway (Figure 5C-D). As a neuropeptide cargo, we used NPY fused to SBP and EGFP, as described previously (Emperador-Melero *et al*, 2018). In the absence of biotin, NPY-EGFP-SBP was diffusely distributed along cell soma and processes, in agreement with its localization to the ER. Addition of biotin induced a wave of NPY-EGFP-SBP accumulating in the Golgi first, and then leaving the Golgi as discrete puncta (Figure 5E, supplemental video 1). These puncta co-localized with an endogenous DCV marker, CHGA, indicating correct packaging of the NPY fusion protein into DCVs during the RUSH assay (Figure 5 supplement 2). Formation of such puncta and clearance of NPY-EGFP-SBP from the Golgi was observed in neurons of both genotypes. We next compared the speed of NPY-EGFP-SBP flux through the Golgi by plotting EGFP intensity within the Golgi area over time (Figure 5F). The peak of fluorescence intensity in the Golgi was reached 5.6 min faster in DKO than in WT, indicating that the speed of Golgi filling is increased in DKO (Figure 5G). The speed of Golgi clearance – determined by fitting a first order exponential decay curve to the data after the peak – was also increased (by 20%) in DKO neurons (Figure 5H). Thus, loss of tomosyns does not impede export of the overexpressed neuropeptide cargo from Golgi but, conversely, accelerates it.

**Figure 5.**
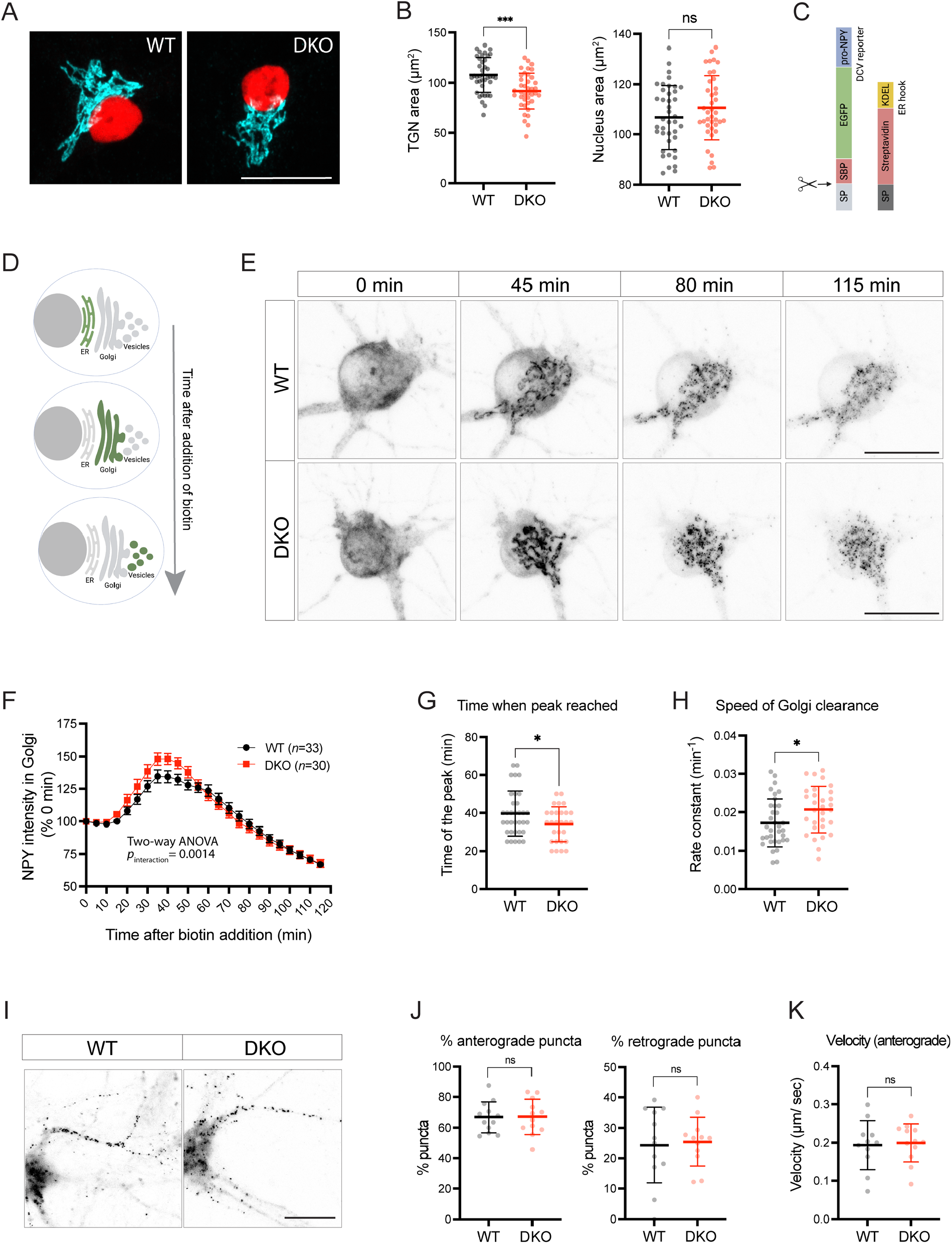
Loss of tomosyns results in the accelerated trafficking of a DCV cargo through Golgi. (A) TGN area is decreased in DKO neurons. Representative images of immunostained TGN marker, TMEM87A (shown in cyan). Nuclei are labeled in red due to expression of mCherry-tagged Cre or ΔCre. Scale bar 20 µm. (B) Quantification of the area of the TGN and nuclei from confocal microscopy images exemplified in (A). Data are shown as mean ± SD and were analyzed using a two-tailed unpaired *t*-test. n=39-40 neurons/ genotype. \*\*\**p* < 0.001; ns: not significant. (C) Design of the construct for the RUSH assay. As a DCV cargo, human pre-pro-NPY fused to SBP and EGFP was used. The cargo was immobilized at the ER due to interaction with the ER hook, consisting of streptavidin fused to the ER-retention/ retrieval signal KDEL. (D) Scheme illustrating synchronized trafficking of a DCV cargo through the secretory pathway upon addition of biotin in the RUSH assay. (E) Snapshots of time-lapse videos taken after addition of biotin in the RUSH assay. Time passed after addition of biotin is shown on top. DCV cargo passes through the Golgi in roughly 45 minutes after addition of biotin and gets concentrated in fine puncta that eventually leave cell soma. Scale bar 20 µm. (F) NPY-EGFP intensity in the Golgi area plotted against time after addition of biotin. Intensity values were normalized to the basal levels (before addition of biotin). Data are shown as mean ± SEM and were analyzed by two-way ANOVA. n=30-34 neurons/ genotype. (G) Time required to reach maximum NPY-EGFP intensity at the Golgi, quantified from (F), is reduced in DKO. Data are shown as mean ± SD and were analyzed by a two-tailed unpaired *t*-test. \**p* < 0.05. (H) Speed of NPY-EGFP export from the Golgi is increased in DKO neurons. Rate constants were calculated from first order decay curves fitted to the data shown in (F). Data are shown as mean ± SD and were analyzed by a two-tailed unpaired *t*-test. \**p* < 0.05. (I) Snapshots of time-lapse videos taken 90 min after addition of biotin in the RUSH assay with the focus on neurites, where newly made DCVs trafficked after exit from the Golgi. (J) Proportion of newly made DCVs trafficking in anterograde and retrograde directions as quantified from the time-lapse videos exemplified in (I). Majority of vesicles trafficked anterogradely in neurons of both genotypes. Data are shown as mean ± SD and were analyzed by a two-tailed unpaired *t*-test. n=12 neurons/ genotype. ns: not significant. (k) Speed of anterogradely trafficking vesicles as quantified from the time-lapse videos exemplified in (I). Data are shown as mean ± SD and were analyzed by a two-tailed unpaired *t*-test. n=12 neurons/ genotype. ns: not significant.

We also traced the fate of newly made RUSH vesicles 90 minutes after addition of biotin. After the exit from Golgi, these vesicles travelled predominantly in one neurite, morphologically identified as the axon (Figure 5I). Most axonal vesicles (67%) traveled in the anterograde direction with a similar speed in neurons of both genotypes (Figure 5J-K), indicating that transport of newly made DCVs is not affected by the loss of tomosyns.

Since the increased rate of NPY trafficking through the Golgi does not explain the reduced levels of DCVs and specific DCV cargos in DKO, we next tested if a more upstream process in DCV production, i.e. expression of DCV genes, is impaired in the absence of tomosyns. To this end, we tested mRNA expression of *Bdnf* (Figure 6A-B), some general DCV markers (Figure 6C), and neuropeptides most abundantly expressed in hippocampal neurons (Figure 6D). Transcription of the *Bdnf* gene yields at least nine transcripts, five of which were readily detectable in cultured hippocampal neurons (Figure 6A). Of those, four were decreased in DKO neurons (Figure 6B). The reduction in expression varied between transcript isoforms. DKO neurons showed the strongest decrease (by 60%) in the isoform 1 mRNA, while levels of the isoform 6 – a much less abundant transcript – were unaffected in DKO. All isoforms encode the same protein, therefore reduced mRNA levels of the two highest expressed isoforms (1 and 4) can explain markedly decreased BDNF levels in DKO neurons.

**Figure 6.**
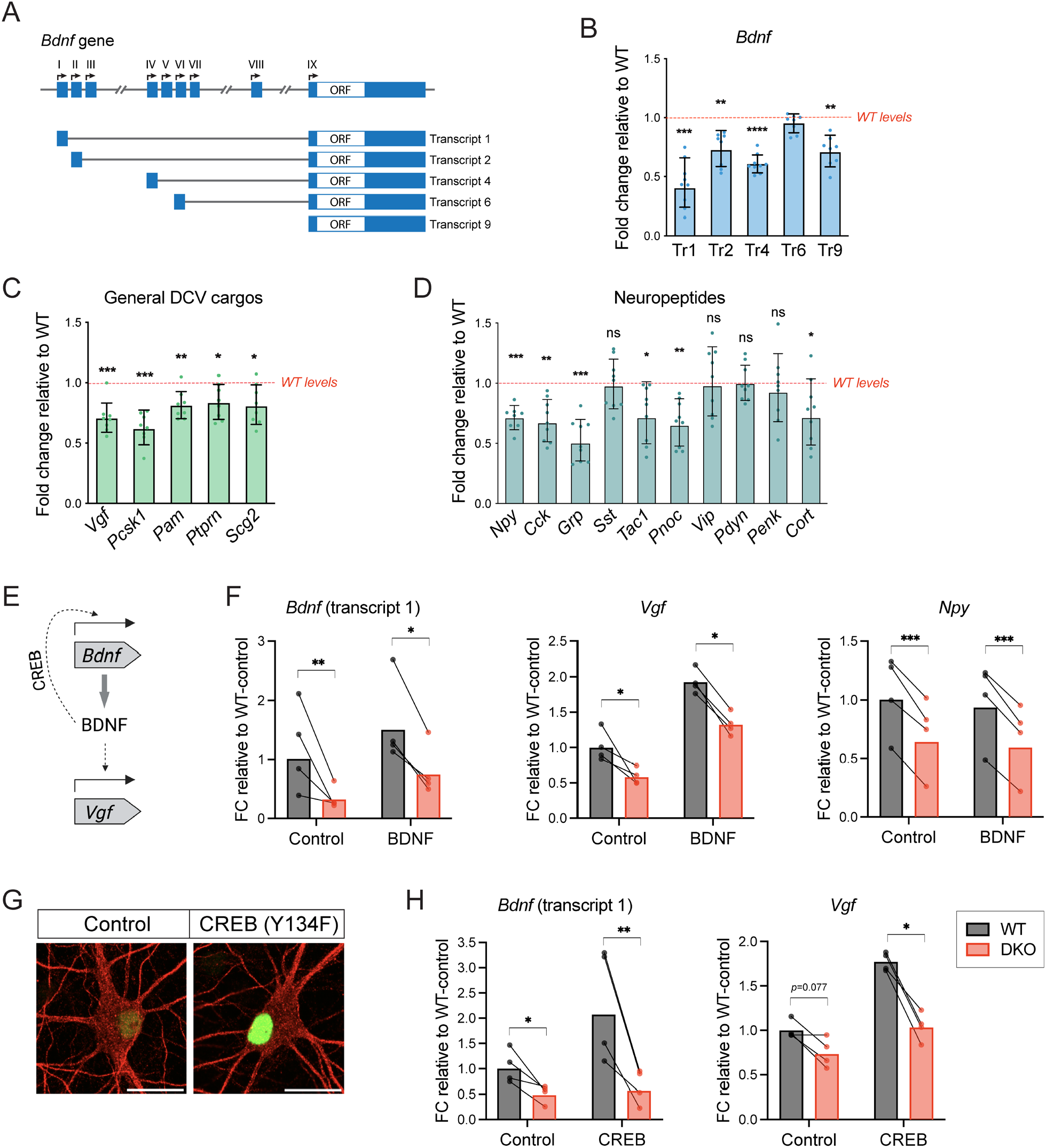
Gene expression of multiple DCV cargos is decreased in DKO neurons. (A) Schematic representation of mouse *Bdnf* gene structure showing nine alternative promoter regions giving rise to multiple transcript isoforms, among which isoforms 1, 2, 4, 6 and 9 are most abundant in the hippocampus. (B) mRNA levels of different *Bdnf* transcript isoforms as assessed by quantitative RT-PCR and normalized to *Gapdh*. Data are shown as fold changes (FC) relative to WT levels in the corresponding culture preparation. n=8 culture preparations/ genotype. Dots represent individual cultures, bar graphs are geometric means, error bars are geometric SD. Log2FC (ΔΔCt) were analyzed by one-sample *t*-test. \*\**p* < 0.01; \*\*\**p* < 0.001. (C) mRNA levels of general DCV cargos as assessed by quantitative RT-PCR and normalized to *Gapdh*. Data are shown and analyzed as described in (B). \**p* < 0.05; \*\**p* < 0.01; \*\*\**p* < 0.001. (D) mRNA levels of most abundant hippocampal neuropeptides as assessed by quantitative RT-PCR and normalized to *Gapdh*. Neuropeptides are sorted by their abundance in hippocampal neurons. Data are shown and analyzed as described in (B). \**p* < 0.05; \*\**p* < 0.01; \*\*\**p* < 0.001, ns: not significant. (E) Scheme depicting a function of BDNF in regulating transcription of its own gene, as well as of other genes (e.g., *Vgf*), which, at least in part, is mediated by the transcription factor CREB. (F) mRNA levels of *Bdnf, Vgf* and *Npy* upon treatment of cultured neurons with recombinant BDNF (50 ng/ml, 3h). The treatment results in an increase of *Bdnf* and *Vgf* mRNA levels in both genotypes (two-way ANOVA *p* _treatment_ < 0.05) but does not rescue the decreased mRNA levels in DKO (red bars). Data are shown as FC relative to untreated WT in the corresponding culture preparation. n=4 culture preparations/ genotype. Dots represent independent culture preparations; bar graphs are geometric means. Log2FC (ΔΔCt) were analyzed by repeated measures two-way ANOVA with post hoc Šidák’s multiple comparisons test. \**p* < 0.05; \*\**p* < 0.01; \*\*\**p* < 0.001. (G) Validation of the expression of CREB (Y134), a constitutively active form of CREB, by ICC in neurons transduced with CREB (Y134)-encoding lentivirus. Immunostaining of CREB phosphorylated at S133 is shown in green, MAP2 is shown in red. Scale bar 20 µm. (H) Overexpression of CREB (Y134) does not rescue reduced mRNA levels of *Bdnf* and *Vgf* in DKO neurons. Data are shown and analyzed as described in (F). \**p* < 0.05; \*\**p* < 0.01.

mRNA expression of all tested general DCV cargos was decreased in DKO as well (Figure 6C), in close agreement with the changes observed in the proteomic analyses (Figure 3). Among neuropeptide genes, *Grp* (gastrin related peptide) and *Pnoc* (nociceptin, nocistatin) showed the strongest reduction in DKO (by 50% and 35%, respectively), while genes encoding cholecystokinin, NPY, tachykinins and cortistatin displayed a reduction by around 30% (Figure 6D). Interestingly, somatostatin (*Sst*), a neuropeptide closely related to cortistatin, was unaltered at the mRNA level in DKO neurons, indicating a certain specificity in expression changes induced by the loss of tomosyns. Based on the single-cell transcriptomics data (Saunders *et al*, 2018), expression of the affected genes was not confined to a specific neuronal population (e. g. glutamatergic or GABAergic neurons). These data suggest that the loss of tomosyns causes a widespread dysregulation in expression of general DCV cargos and specific neuropeptides across different subpopulations of hippocampal neurons.

Since BDNF regulates expression of many secretory genes, including its own gene, we tested whether the decreased mRNA levels of *Bdnf* and its known targets (*Vgf, Npy*) is a consequence of a potentially decreased BDNF secretion (Figure 6E). We analyzed expression of *Bdnf*, Vgf and *Npy* after treatment of neurons with recombinant BDNF (50 ng/ml for 3 h). Though this treatment increased mRNA levels of *Bdnf* (isoform 1) and *Vgf* in neurons of both genotypes, it did not rescue the impaired expression of these genes in DKO neurons (Figure 6F). *Npy* expression did not respond to the BDNF treatment. Thus, decreased BDNF levels are not the cause of the impaired DCV gene expression in the absence of tomosyns. Transcription factor CREB is one of the few established regulators of neuronal DCV gene expression in response to many factors, including BDNF and neuronal activity (Moore *et al*, 1996; Finkbeiner *et al*, 1997; Tao *et al*, 1998). We next tested whether overexpression of a constitutively active form of CREB – carrying the Y134F mutation that lowers the threshold for the activating S133 phosphorylation (Du *et al*, 2000) – rescues the transcriptional defect of DKO neurons. This intervention failed to rescue expression of *Bdnf* and *Vgf* in DKO neurons (Figure 6G-H). In addition, levels of activated CREB (phosphorylated on S133) were not affected in DKO neurons as evidenced by WB on neuronal lysates using a phospho-specific antibody (Figure 6 supplement 1). These data suggest that altered CREB activity does not account for the reduced expression of *Bdnf* and other DCV cargo genes in DKO neurons.

## Discussion

In this study, we examined the role of tomosyns in exocytosis of neuropeptide containing DCVs in mammalian neurons. Using pHluorin-based DCV fusion reporter, we found that the loss of tomosyns does not alter DCV exocytosis but lowers intracellular levels of the DCV reporter. We corroborated the latter finding by showing that levels of many endogenous DCV cargos are decreased in tomosyn DKO neurons by three independent approaches (ICC, WB, and mass spectrometry). The effect on DCV pool and/or composition was accompanied by a smaller DCV size, increased speed of the DCV reporter flux through the TGN, and decreased mRNA levels of many (but not all) DCV cargos. Taken together, these data suggest that tomosyn is dispensable for the control of DCV fusion, but, paradoxically, has an impact on the expression of DCV genes and intracellular neuropeptide levels.

The lack of the effect on neuronal DCV fusion is surprising, given that tomosyn regulates SV release in neurons and secretory granule exocytosis in neuroendocrine cells and platelets. However, phenotypes caused by the loss of tomosyn expression in different cell types are discrepant. While loss of tomosyn resulted in enhanced SV fusion in the mouse brain and invertebrate nervous system (Sakisaka *et al*, 2008; Ben-Simon *et al*, 2015; Gracheva *et al*, 2006; McEwen *et al*, 2006; Chen *et al*, 2011; Sauvola *et al*, 2021), tomosyn deficiency in platelets, INS-1E cells and rat superior cervical ganglion neurons led to an opposite phenotype – a strong reduction in secretion (Ye *et al*, 2014; Zhu *et al*, 2014; Cheviet *et al*, 2006; Baba *et al*, 2005). No unifying model explains the discrepant effects of tomosyn on vesicle fusion.

DCVs and SVs differ in release properties, and recent studies shed light on some components of the molecular machinery contributing to these differences (Persoon *et al*, 2019; Moro *et al*, 2021). In contrast to SVs, which are released in response to a single electric pulse, neuronal DCVs require prolonged high-frequency stimulation for induction of exocytosis (Hartmann *et al*, 2001; Xia *et al*, 2009; Persoon *et al*, 2018). Several factors can explain the reluctance of DCV to fuse, among them the loose coupling of DCVs to the sites of calcium entry, requiring an increase in calcium levels reaching beyond an immediate radius of a calcium channel (Mansvelder & Kits, 2000). Interestingly, the inhibitory effect of tomosyn on DCV fusion in adrenal chromaffin cells is calcium-dependent and could be overridden by an increase in intracellular calcium (Yizhar *et al*, 2004). Thus, it is possible that under high-frequency stimulation required for the induction of neuronal DCV exocytosis, the regulatory effect of tomosyn on vesicle fusion is lost.

Altogether, the discrepant effects of tomosyn on fusion in different cell types suggest that tomosyn does not act solely as an inhibitor of fusogenic SNARE complex formation. In line with this, loss of tomosyns in primary neurons did not alter the amount of assembled ternary SDS-resistant SNARE complexes (Figure 1H). In addition to the SNARE domain, tomosyn contains multiple WD40-repeats folded into two bulky beta-propellers. The two propellers are, in fact, the largest and evolutionary best conserved part of tomosyn. Several interaction partners of tomosyn, such as synapsin and Rab3, bind to the propeller domain (Cazares *et al*, 2016). These interactions may mediate targeting of tomosyn to SV membrane (Takamori *et al*, 2006) and act in tomosyn-dependent regulation of SV pools (Cazares *et al*, 2016). Interestingly, yeast orthologs of tomosyn, SRO7 and SRO77, localize to post-Golgi vesicles and act as vesicle tethers at the plasma membrane via interaction with the RAB3 ortholog Sec4 GTPase (Lehman *et al*, 1999; Rossi *et al*, 2018). Thus, tomosyn may affect steps far upstream of vesicle fusion via WD40-domain mediated interactions. In this case, the effect of tomosyn on exocytosis of different types of vesicles would depend on the mechanisms of vesicle tethering/ docking, which differ between SVs and DCVs. Tomosyn co-purifies with DCVs in chromaffin cells (Wegrzyn *et al*, 2010) and travels together with DCVs in neurons (Geerts *et al*, 2017) but its loss did not affect neuronal DCV distribution and trafficking (Figure 2A-D, 5J-K), suggesting that tomosyn is dispensable for DCV tethering in neurons.

Despite the lack of effect on DCV exocytosis, loss of tomosyns resulted in reduced intracellular levels of many DCV cargos. The latter phenotype is consistent with the effect of tomosyn in platelets, where its loss led to a strong decrease in many secretory cargos, despite the fact that the release of these cargos was reduced, not enhanced (Ye *et al*, 2014). Similarly, loss of SRO7/77 in yeast resulted in the accumulation of secretory vesicles with an altered composition (Forsmark *et al*, 2011). Taken together, these data suggest a role of tomosyn and its orthologs in the formation of secretory granules. The accumulation of tomosyn at the TGN (Figure 5 supplement 1) and decreased DCV size in DKO neurons (Figure 4A-C) are consistent with such a role. However, loss of tomosyns did not impair formation of new NPY vesicles in the RUSH assay, indicating that tomosyns are dispensable for DCV biogenesis (Figure 5E, supplemental video 1). Interestingly, NPY flux through the Golgi was slightly accelerated in the absence of tomosyns (Figure 5F-H), which, in combination with the decreased TGN size in DKO, suggests a role of tomosyn(s) in the TGN membrane trafficking. Knockdown of tomosyn in hippocampal neurons was shown to modulate activity of RhoA GTPase (Shen *et al*, 2020), which, in turn, regulates membrane trafficking at the Golgi (Quassollo *et al*, 2015). On the other hand, the decreased TGN size can be explained by the reduced expression of many secretory cargos in DKO neurons, since Golgi size is regulated by the secretory cargo load (Sengupta & Linstedt, 2011).

The most unexpected observation of this study is a decrease in mRNA expression of many DCV cargos in DKO neurons (Figure 6B-D), suggesting a role of tomosyn(s) in regulation of the DCV gene transcription. We ruled out the simplest mechanism of such regulation, whereby decreased BDNF levels in DKO would result in an impaired transcriptional feedback loop.

Treatment of DKO neurons with BDNF, as well as overexpression of the activated CREB that mediates this autoregulation (Esvald *et al*, 2020), did not rescue decreased *Bdnf* and *Vgf* mRNA levels (Figure 6F-H). Considering the established effect of tomosyn on SV exocytosis and synaptic activity, it is possible that altered neuronal firing patterns contribute to the decreased expression of DCV genes in DKO. *Bdnf* and many neuropeptide genes are known targets of activity-dependent signal transduction (Zafra *et al*, 1990; Ernfors *et al*, 1991; Isackson *et al*, 1991; MacArthur & Eiden, 1996). If indeed caused by chronically altered firing patterns, the transcriptional defect of DKO neurons is cell-autonomous, since the decrease in BDNF was detected in both single neuron and network cultures. In addition to the decreased mRNA expression of DCV genes, other factors likely contribute to the reduction of DCV cargos in DKO neurons, since loss of tomosyns also affected levels of the DCV fusion reporter expressed under control of the synapsin promoter, even after silencing of neuronal activity with TTX and despite the normal activity of this promoter in DKO (Figure 1H-I, Figure 1 supplement 3).

In summary, our current findings indicate that tomosyns are dispensable for neuronal DCV exocytosis but regulate intracellular neuropeptide levels. Decreased levels of many neuropeptides and BDNF at the critical moment of neuronal development may promote the onset of neurodevelopmental disorders associated with mutations in *STXBP5/5L* genes.

## Materials and methods

### Mice

*Stxbp5*^lox/lox^, *Stxbp5l*^lox/lox^ double conditional null mice were generated by breeding two lines with the individually targeted *Stxbp5* and *Stxbp5l* alleles. To generate *Stxbp5*^lox/+^ mice, *lox2272* recombination sites flanking exon 2 of *Stxbp5* (98 bp in size) were introduced into the genome via homologous recombination in C57Bl/6 embryonic stem (ES) cells (Cyagen Animal Model Services, Santa Clara, USA). Lox2272 recombines efficiently with other lox2272 sites, but not with loxP (Lee & Saito, 1998). Deletion of exon 2 causes a frameshift in all tomosyn reference transcripts (NM_001081344.3, NM_001408063.1, NM_001408064.1 and NM_001408065.1). The targeting vector contained a neomycin resistance marker, flanked by self-deleting anchor sites and placed between exon 2 and the lox2272 site in intron 2. Targeted ES clones were injected into blastocysts to produce germ-line chimeras, which were mated to C57Bl/6J wild-type mice. *Stxbp5* line genotyping was performed using the following primers (5’ ACTTAGCGCGGAGGGTTTTGTC 3’ and 5’ CGTAGGCTTTTAAATCACCGCTGT 3’) to amplify *Stxbp5*^+^ or S*txbp5*^lox^ specific products of 196 and 236 bp in size, respectively. The *Stxbp5l*^lox/+^ line was generated independently as described in (Geerts *et al*, 2015). In these mice, the paralogous exon 3 (also 98 bp in size) is flanked by loxP sites. Cre-mediated excision induces a frameshift in all reference transcripts (NM_172440.3, NM_001114611.1, NM_001114612.1, NM_001114613.1, also named xb-, s-, b-and m-tomosyn-2, respectively). *Stxbp5l* line genotyping was performed for *Stxbp5l* as described (Geerts *et al*, 2015). After interbreeding the two mouse lines, both backcrossed to an inbred C57Bl/6J genetic background, newborn pups from homozygous *Stxbp5*^lox/lox^, *Stxbp5l*^lox/lox^ matings were used for the preparation of neuronal cultures in all the described experiments. Mice were housed and used for experiments according to institutional guidelines and Dutch laws.

### Neuronal cultures

Mouse hippocampal cultures were prepared from newborn pups. Normally, hippocampi from several pups from one nest were pooled in one culture preparation, which was considered as one biological replicate. Hippocampi were dissected in Hanks Balanced Salt Solution (HBSS, Sigma H9394) supplemented with 10 mM HEPES (pH 7.2-7.5, Gibco™ 15630056) and digested with 0.25% trypsin (Gibco™ 15090046) at 37°C for 15 min. After the digestion, tissue pieces were washed in HBSS and triturated using fire-polished Pasteur pipettes in Dulbecco’s modified Eagle medium (DMEM, VWR Life Science VWRC392-0415) supplemented with 10% fetal calf serum (FCS, Gibco™ 10270), non-essential amino acids (Sigma M7145), and antibiotics (penicillin/ streptomycin, Gibco™ 15140122). For most immunocytochemistry (ICC) experiments, cell suspension was sparsely plated on a feeder layer of rat glia at a density of 20-30*10^3^/ well on 15 mm coverslips. For BDNF ICC and pHluorin imaging experiments, neurons were plated on glial micro-islands at a density of 1500/ well (to obtain single neuron cultures). For RUSH, rat glia were omitted, and neurons were plated on glass coverslips at a density of approximately 120*10^3^/ well on 15 mm coverslips. Prior to plating, glass coverslips were coated with poly-L-ornithine (Sigma P4957) and laminin (Sigma L2020). For western blot, proteomics, electron microscopy and qPCR experiments, neurons were plated on coated plastic without the glia feeder layer at a density of 400*10^3^/ well in a 6-well plate. Cultures were maintained in a humified incubator at 37 °C and 5% CO_2_ in Neurobasal medium (Gibco™ 21103049) supplemented with B27 (Gibco™ 17504044), 10 mM HEPES, GlutaMAX™ (Gibco™ 35050038) and antibiotics (penicillin/ streptomycin, Gibco™ 15140122). To induce loss of tomosyns, cultures were transduced within 8 h after plating with a lentivirus encoding Cre-recombinase expressed under control of synapsin promoter (pSyn-Cre-mCherry). As control, neurons from the same culture preparation were transduced with *Cre*-recombinase lacking the DNA-binding domain (pSyn-ΔCre-mCherry). Experiments were performed, if not otherwise stated, on day *in vitro* (DIV) 14.

Rat glia were prepared from cortices of newborn rats. Cortices were digested in papain (Worthington Biochemical Corporation LS003127) at 37°C for 45 min and triturated in DMEM supplemented with 10% FCS, non-essential amino acids, and antibiotics. Glia were plated and expanded in T175 flasks. For making single neuron cultures, glia were plated on micro-islands of growth permissive substrate (mix of collagen I and poly-D-lysine, Corning 354236 and Sigma P6407, respectively) that were printed on a layer of 0.15% agarose.

### Visualization and analysis of DCV fusion

Single neuron cultures were transduced on DIV8 with a lentivirus encoding the N-terminal part of human NPY fused to super-ecliptic pHluorin (Figure 1 supplement 2). DIV14-15 neurons were used for live imaging on a Nikon Ti Eclipse inverted microscope equipped with A1R confocal module and Andor DU-897 camera (with 512×512 pixels frame) under 40x oil objective (numerical aperture, NA 1.3). LU4A laser unit with 488 nm wavelength laser was used as the illumination source. Coverslips were transferred in an imaging chamber in Tyrode’s solution (119 mM NaCl, 2.5 mM KCl, 2 mM CaCl_2_, 2 mM MgCl_2_, 25 mM HEPES and 30 mM Glucose, pH 7.4, 280 mOsmol) and time-lapse videos were recorded at 2 Hz frequency for 2 min at room temperature under constant perfusion with Tyrode’s. After 30 sec of baseline recording, neurons were subjected to electric field stimulation (16 trains, each consisting of 50 pulses at 50 Hz and separated by 500 msec intervals) delivered by platinum electrodes. At the end of each recording, 50 mM NH_4_Cl-containing Tyrode’s solution (with NaCl concentration reduced to 69 mM to maintain osmolarity) was applied to visualize the intracellular pool of DCVs. Analysis of the time-lapse recordings was performed as described (Persoon *et al*, 2019). In short, fusion events were detected by a sudden rise in fluorescence intensity. 3×3 pixel regions of interest (ROI) were placed on time-lapse recordings using a custom-made script in Fiji (Schindelin *et al*, 2012). Change in fluorescence intensity (F) within these ROIs was plotted as ΔF/F_0_, where F_0_ is an average of the first 10 frames of acquisition. Resulting traces were checked using a custom-made script in MATLAB, and only events with F/F_0_ ≥ 2 SD and rise time < 1 sec were counted. Total number of intracellular vesicles (intracellular pool) was determined as the number of fluorescent puncta after the ammonium pulse corrected to account for overlapping puncta. Released fraction was calculated as the number of fusion events per neuron divided by the intracellular pool of DCVs.

### Immunocytochemistry

Neurons grown on coverslips were fixed by application of phosphate-buffered saline (PBS) containing 4% paraformaldehyde (PFA, Sigma P6148) and 4% sucrose for 10 min. After fixation, neurons were washed and the residual PFA was quenched by application of 0.1 M glycine for 5 min. Neurons were permeabilized and blocked in PBS containing 0.1% Triton™ X-100 (Fisher Chemical T/3751/08) and 2% normal goat serum (Gibco™ 16210-072) (‘blocking buffer’) for 30 min. Primary antibodies diluted in blocking buffer were applied for 1 h at room temperature. Primary antibodies used in this study and their dilutions are listed in Table 1. After washing with PBS, neurons were incubated with Alexa Fluor-coupled secondary antibodies diluted at 1:500 for 45 min. Stained coverslips were mounted in Mowiol.

**Table 1.** List of primary antibodies used in this study.

| Antigen | Catalogue number | Type of experiment | Dilution |
| --- | --- | --- | --- |
| Syntaxin 1 | Sigma S0664 | WB | 1:2000 |
| Tomosyn | Synaptic Systems 183103 | WB | 1:1000 |
|  |  | ICC | 1:500 |
| Tomosyn-2 | Synaptic Systems 183203 | WB | 1:1000 |
| alpha-Tubulin | Synaptic Systems 302211 | WB | 1:1000 |
| Actin | Merck MAB1501 | WB | 1:1000 |
| Synaptophysin | Synaptic Systems 101004 | ICC | 1:1000 |
| MAP2 | Abcam ab5392 | ICC | 1:5000 |
| IA-2 (PTPRN) | Merck MABS469 | ICC, WB | 1:100 |
| CHGA | Synaptic Systems 259003 | ICC | 1:500 |
| CHGB | Synaptic Systems 259103 | ICC | 1:500 |
| BDNF | DSHB hybridoma product BDNF #9 (supernatant) * | ICC | 1:4 |
| LAMP1 | DSHB hybridoma product 1D4B (supernatant) ** | ICC | 1:20 |
|  |  | WB | 1:200 |
| LIMP2 | Novus Biologicals NB400-129 | WB | 1:1000 |
| TMEM87A | Novus Biologicals NBP1-90532 | ICC | 1:100 |
| TGN38 | Bio-Rad AHP499G | ICC | 1:100 |
| pCREB (S133) | R&D Systems MAB6906 | ICC | 1:50 |
|  |  | WB | 1:1000 |

\* This monoclonal antibody developed by Barde Y.-A. was obtained from the Developmental Studies Hybridoma Bank, created by the NICHD of the NIH and maintained at The University of Iowa, Department of Biology, Iowa City, IA 52242.
\*\* This monoclonal antibody developed by August J.T. was obtained from the Developmental Studies Hybridoma Bank, created by the NICHD of the NIH and maintained at The University of Iowa, Department of Biology, Iowa City, IA 52242.

### Confocal microscopy of fixed neurons

Imaging of stained neurons was performed on a Nikon Ti Eclipse inverted microscope equipped with A1R confocal module and LU4A laser unit. Images were acquired under 40x (NA 1.3) or 60x (NA 1.4) oil objectives in the confocal mode. Acquisition parameters (zoom, image size, scanning speed, laser power and gain) were set using NIS software to achieve appropriate image resolution and avoid signal saturation. Three to five z-sections with an interval of 0.2 µm were collected per image.

Analysis of staining intensity was performed on maximum intensity projection of z-stacks using a custom-made macro in Fiji, as described (Moro *et al*, 2020). Masks of the neuronal somata and neurites were obtained based on MAP2 signal. Puncta of DCV cargos (BDNF, IA-2) were detected within the neurite mask based on their intensity and dimensions.

### SDS-PAGE western blot (WB) analysis

To analyze protein levels, high-density neuronal cultures (400*10^3^/well in a 6-well plate) were lysed in Laemmli sample buffer. Lysates were heated at 95°Cfor 5 min, separated by SDS-PAGE using either home-made tris-glycine or commercially available Mini-PROTEAN TGX Stain-Free gels (Bio-Rad 4568096), and transferred to nitrocellulose membrane (Bio-Rad 1620115) using wet tank transfer method. Membranes were blocked in 5% milk powder dissolved in Tris-buffered saline containing 0.1% Tween^Ⓡ^ 20 (TBS-T). Membranes were incubated with primary antibodies diluted in TBS-T on a shaking platform overnight at 4°C. Primary antibodies used in this study and their dilutions are listed in Table 1. Horseradish peroxidase coupled secondary antibodies were applied at 1:10000 dilution for 1 h at room temperature. Chemiluminescence-based detection was performed using SuperSignal West Femto Maximum Sensitivity Substrate (Thermo Scientific 34095) on the Odissey Fc imaging system (LI-COR Bioscience). Signal intensities of bands of interest were analyzed using Image Studio Lite Software and normalized to the intensity of a loading control (actin or tubulin). Actin levels were not affected by the loss of tomosyn, as evidenced by comparing band intensities of actin to total protein stain visualized on Mini-PROTEAN TGX Stain-Free gels.

### Detection of SNARE complexes

Detection of SNARE complexes was performed as described previously (Hayashi *et al*, 1994; Otto *et al*, 1997). The protocol is based on the observation that assembled synaptic SNARE complexes (consisting of syntaxin-1, SNAP25 and VAMP2) are preserved in SDS-containing lysis buffer (‘SDS-resistant’) but sensitive to temperatures higher than 60°C. To detect syntaxin-1 in SNARE complexes, high-density neuronal cultures (400K/well in a 6-well plate) were lysed in 1% SDS-containing Laemmli sample buffer, passed through insulin syringe and split in two parts. The first part of the lysates was heated at 95°C for 5 min (for the detection of total syntaxin-1), while the second part was heated at 37°C for 5 min (for the detection of syntaxin-1 in SNARE complexes). Lysates were separated by SDS-PAGE using 4-20% gradient gels (Mini-PROTEAN TGX Stain-Free gels, Biorad #4568096). Protein transfer and detection were performed as described above.

### Proteome analysis by mass spectrometry

DIV12-13 hippocampal neurons grown in high density cultures (400*10^3^/ well in a 6-well plate) were washed once with pre-warmed PBS and lysed in Laemmli sample buffer (50 μl/ well). Lysates were passed through insulin syringe and stored at -80°C until further processing. In-gel digestion with Trypsin/Lys-C Mix solution (Promega) was performed as previously described (Gonzalez-Lozano & Koopmans, 2019). Peptides were analyzed by micro LC-MS/MS using a TripleTOF 5600 mass spectrometer (Sciex, Framingham, MA, USA). The peptides were fractionated with a linear gradient of acetonitrile using a 200 mm Alltima C18 column (300 μm i.d., 3 μm particle size) on an Ultimate 3000 LC system (Dionex, Thermo Scientific, Waltham, MA, USA). Data-independent acquisition was used with Sequential Window Acquisition of all THeoretical mass spectra (SWATH) windows of 8 Da, as previously described (Gonzalez-Lozano *et al*, 2021).

SWATH data was analyzed using DIA-NN (v1.8) (Demichev *et al*, 2020). A spectral library was generated from the complete mouse proteome with a precursor m/z range 430-790. Data was searched with 20 ppm mass accuracy, match-between-runs algorithm (MBR) enabled and robust LC as quantification strategy. Propionamide was selected as fixed modification. Downstream analysis was performed using MS-DAP (v1.0, https://github.com/ftwkoopmans/msdap). The experimental replicate with the lowest number of identified peptides was removed from the analysis (for both genotypes). Only peptides identified and quantified in at least four out of five remaining replicates per genotype were included in the analysis. Normalization was achieved using variance stabilization normalization (Vsn) and mode-between protein methods. The Msqrob algorithm was selected for differential expression analysis, using an FDR adjusted p-value threshold of 0.01 and log2 fold change of 0.3 to discriminate significantly regulated proteins. All data are available in PRIDE repository with the identifier PXD038442. Gene ontology enrichment analysis of the differentially expressed proteins was performed using ShinyGO application v0.75 with an FDR adjusted p-value threshold of 0.05 (http://bioinformatics.sdstate.edu/go/) (Ge *et al*, 2020).

### Electron microscopy

Neurons grown on coated glass coverslips were fixed on DIV14 in 2.5% glutaraldehyde in 0.1M cacodylate buffer. The cells were postfixed in 1% osmium/1% ruthenium and subsequently dehydrated by increasing ethanol concentrations (30%, 50%, 70%, 90%, 96% and 100%) before embedding in EPON. After polymerization of the resin for 72 h at 65 °C, glass coverslips were removed by heating the samples in boiling water. Neuron-rich regions were selected by light microscopy, cut out and mounted on pre-polymerized EPON cylinders. Ultrathin sections of 70-90 nm were cut on an ultra-microtome (Reichert-Jung, Ultracut E) and collected on formvar-coated single slot grids. Finally, the sections were contrasted with uranyl acetate and lead citrate in an ultra-stainer (Leica EM AC20) and imaged in a JEOL1010 transmission electron microscope (JEOL) at 60 kV while being blinded for the experimental conditions. Synapses, somas and DCV-rich areas were photographed by a side-mounted Modera camera (EMSIS GmbH). While remaining blinded for experimental conditions, DCV diameters were measured in iTEM software (Olympus) and synapse parameters were quantified in a custom-written software running in Matlab (Mathworks).

### RUSH assay

The protocol is modified from (Boncompain *et al*, 2012). Neurons were grown in standard neuronal medium except Neurobasal without phenol red (Gibco™ 12348017) was used to minimize an autofluorescent background during live imaging. On DIV7, half of the medium was refreshed with the same medium lacking B27 supplement (to reduce extracellular biotin levels). On DIV9, neurons were transfected using the calcium phosphate transfection method with a plasmid encoding a DCV reporter and an ER hook (pCMV-Streptavidin-KDEL-IRES-SBP-EGFP-NPY) together with a ‘filler’ plasmid (pCMV-mCherry). Neurons were imaged live approximately 16h after the transfection upon addition of 40 μM biotin to the conditioned medium. Time-lapse imaging was performed on a Nikon Ti Eclipse inverted microscope equipped with A1R confocal module under 40x oil objective (NA 1.3). LU4A laser unit with 488 nm wavelength laser was used as the illumination source. Neurons were picked for imaging based on the mCherry fluorescence and were imaged at 37°C and 5% CO_2_ for 2 h after addition of biotin, at a frequency of 1 frame/ 5 min.

Movies were analyzed in Fiji. Possible lateral drift during recordings was corrected using the Image Stabilizer plugin. Mean intensity of NPY-EGFP within the Golgi mask was measured across the frames and plotted against time in GraphPad Prism. The rate constant of Golgi clearance was determined by fitting a one phase exponential decay curve to the decay phase of the intensity plot.

For imaging of DCV trafficking in neurites, transfected neurons were imaged in 80 min after addition of biotin with 0.5 Hz frequency (3 min recordings). Kymographs (space/time plots) were generated from time-lapse videos in Fiji using the KymoResliceWide plugin. Kymographs were analyzed using Kymobutler software (Jakobs *et al*, 2019).

### Quantitative RT-PCR (qRT-PCR)

RNA extraction from neuronal cultures was performed using ISOLATE II RNA Micro Kit (Meridian Bioscience BIO-52073). cDNA was synthesized from purified RNA using SensiFast cDNA Synthesis Kit (Meridian Bioscience BIO-65054). qRT-PCRs were run on a QuantStudio 5 system (ThermoFisher Scientific) using SensiFast SYBR Lo-ROX Kit (Meridian Bioscience BIO-94020). qRT-PCR primers (listed in Table 2) were validated for the efficiency by running qRT-PCRs on 10-fold serial dilutions of cDNA, and for the specificity estimated from melt curves. Only primers with an efficiency of 90-105% were used for experiments. Fold changes (FC) in gene expression were determined using the cycle threshold (CT) comparative method (2^−ddCT^) using *Gapdh* as a housekeeping gene. *Gapdh* CT values were not affected by the genotype or treatment applied. Statistical analysis of the data was performed on log-transformed data (logFC, ddCT).

**Table 2.**
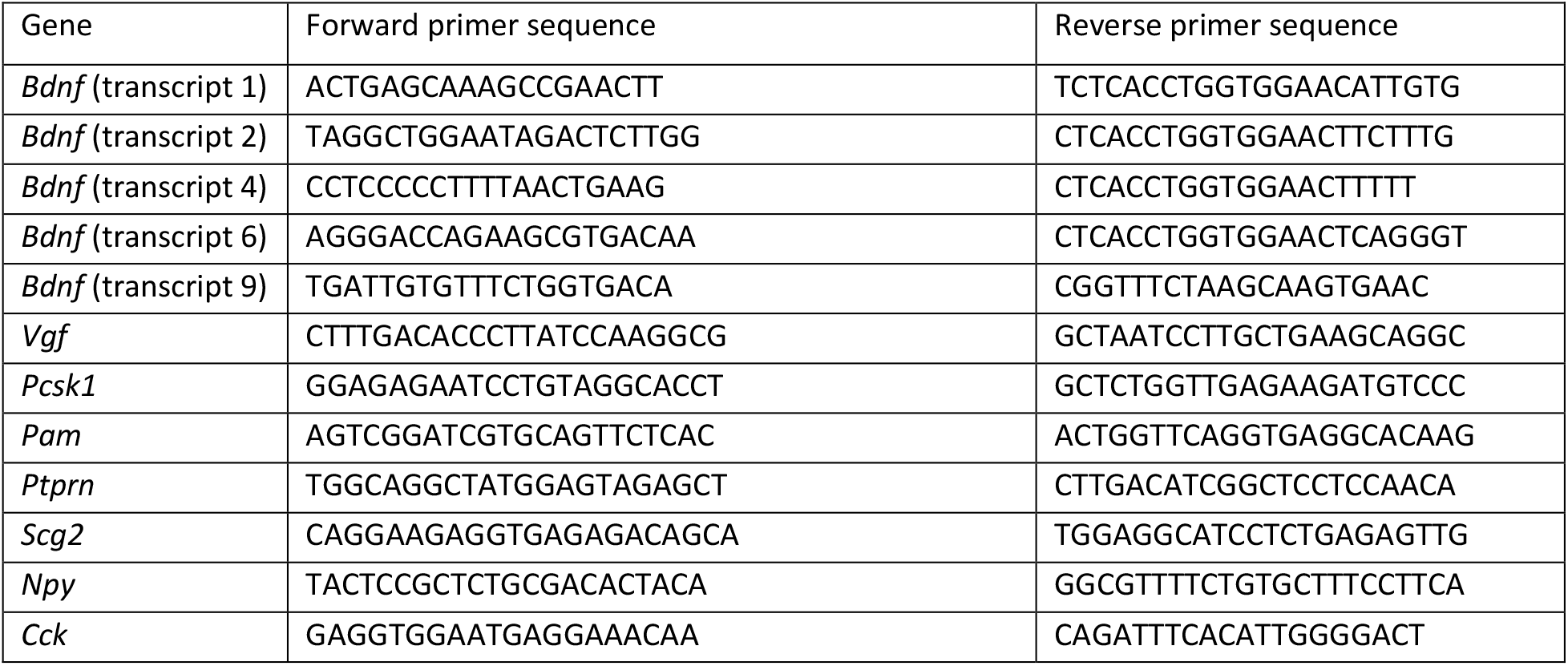
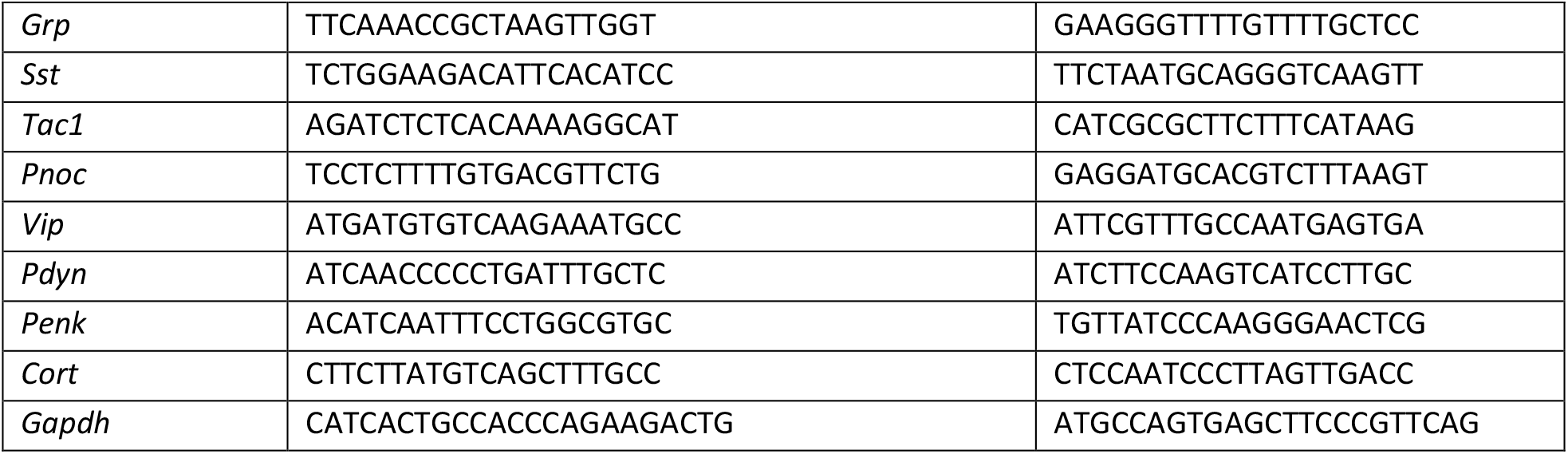
List of qRT-PCR primers used in this study

### Statistical analyses

Statistical analyses were performed using GraphPad Prism software. Data distribution was tested for normality using the D’Agostino-Pearson test. Possible outliers were identified using the ROUT method. An F test was used to compare variances between the groups. Statistical tests used in individual experiments are specified in the corresponding figure legends. Full statistical information, including exact p-values, is provided in Table 3.

**Table 3.**
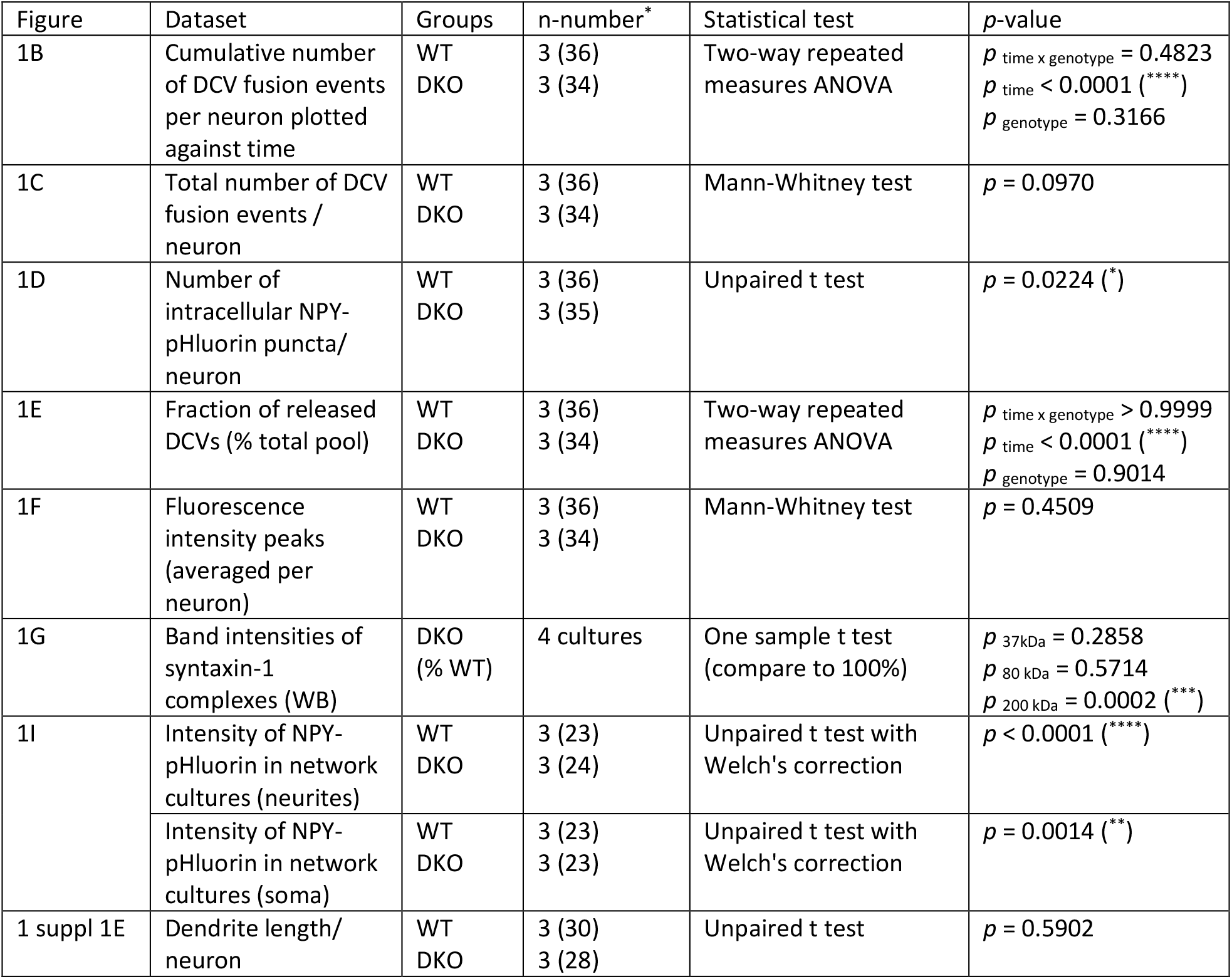
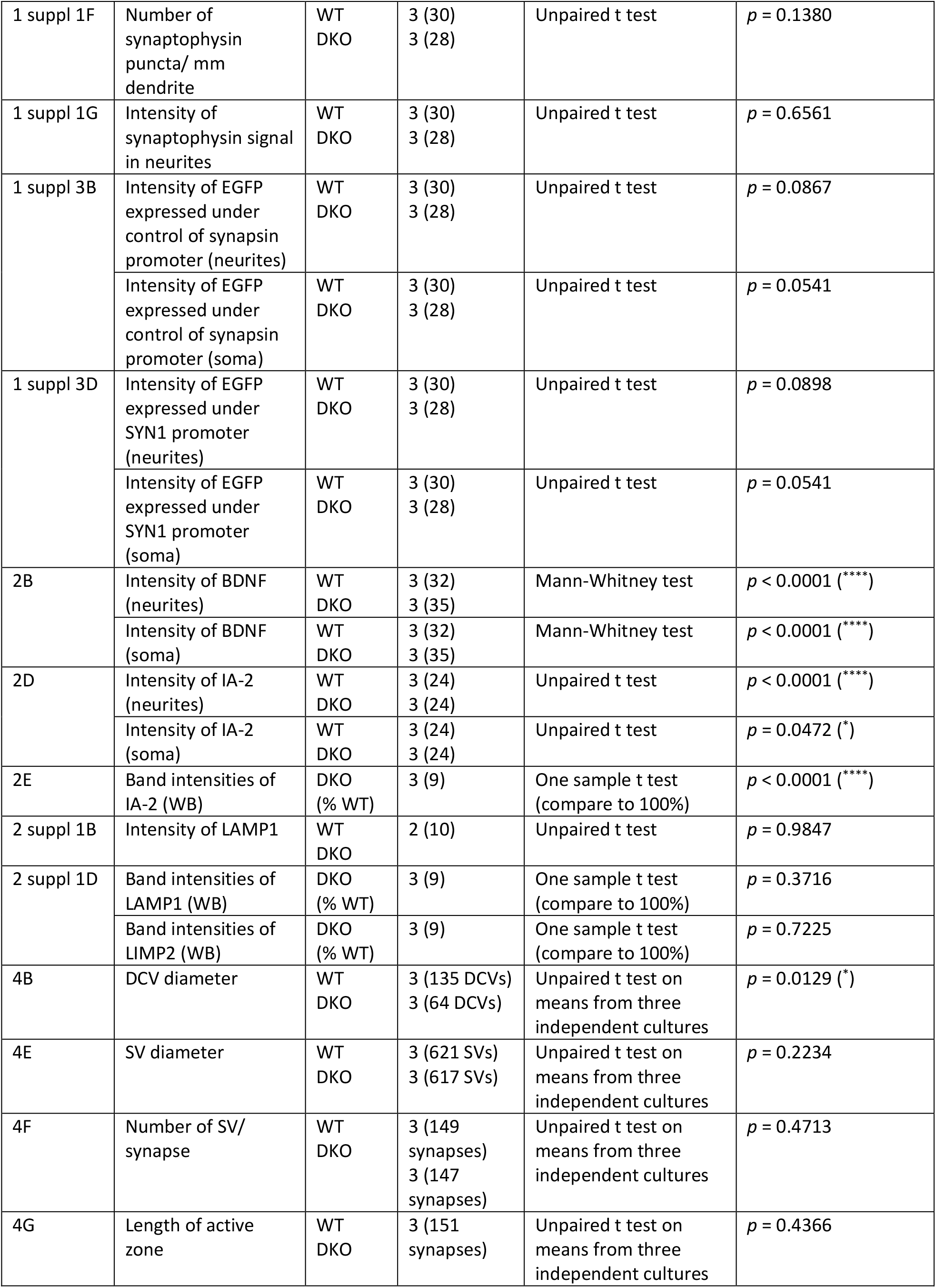
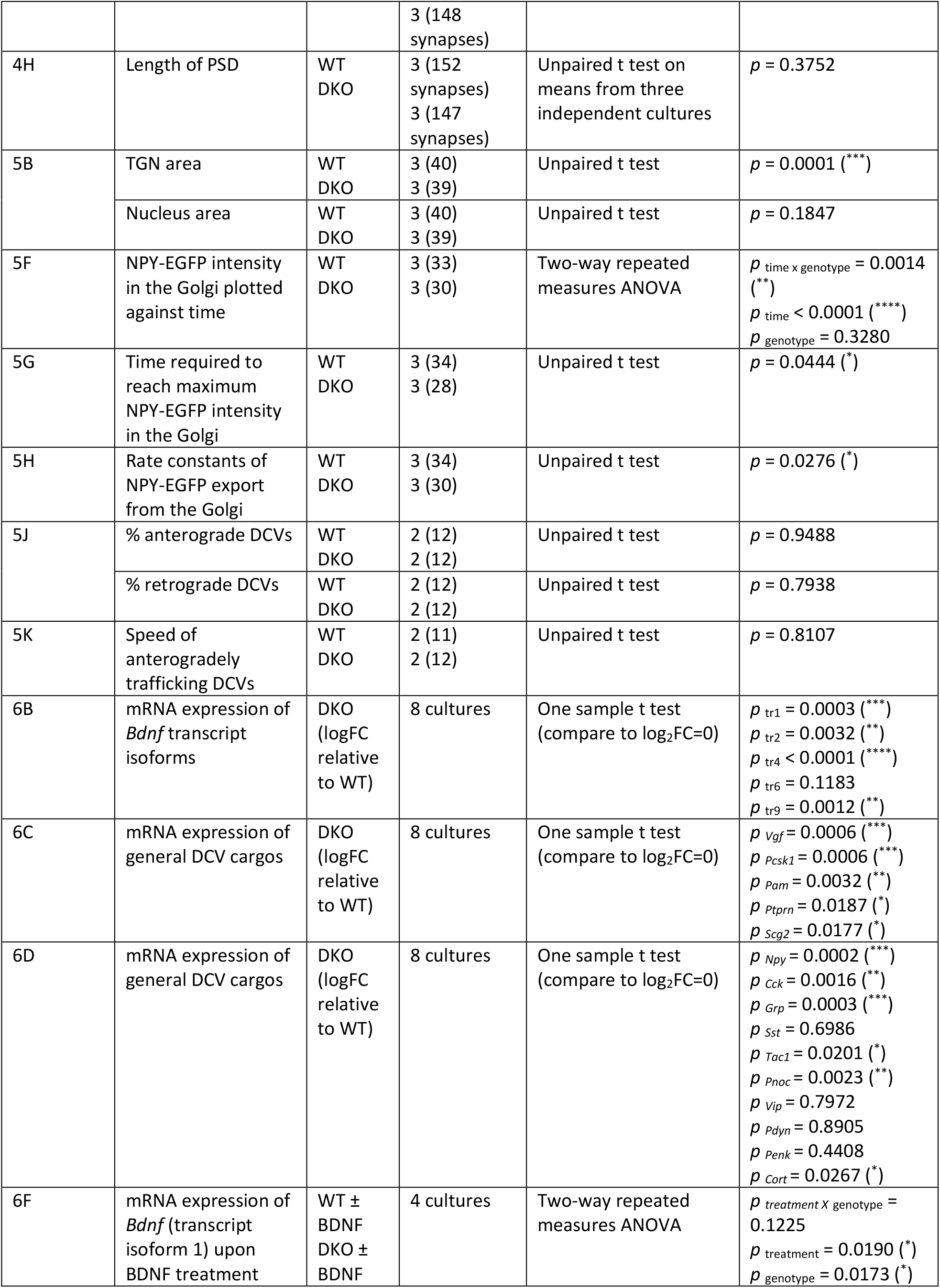
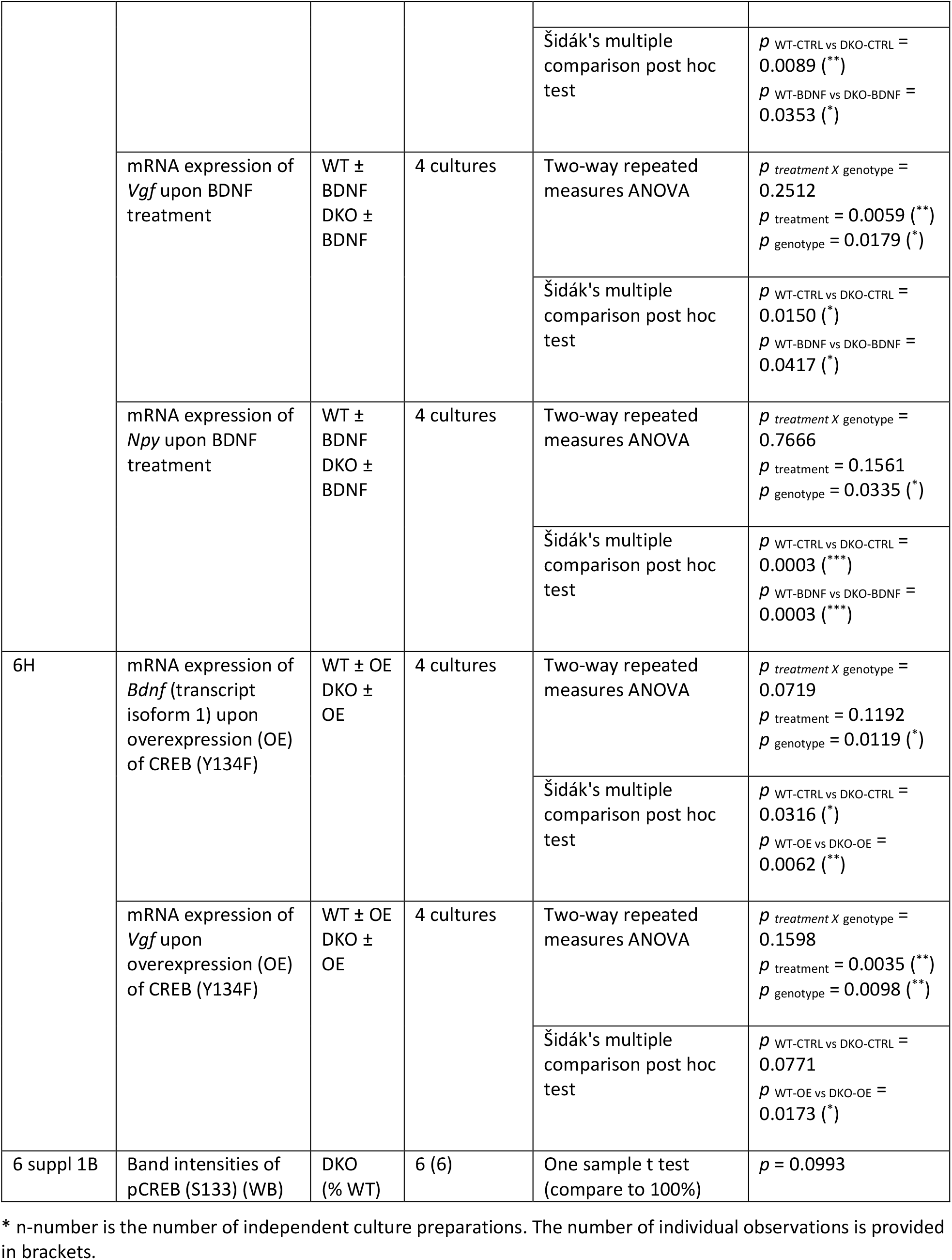
Summary of statistical analyses applied in this study

## Acknowledgements

We are indebted to D. Schut, L. Laan, I. Saarlos, R. Zalm, J. Wortel, J. Hoetjes, R. Dekker, I. Paliukhovich for their excellent technical assistance. This work was supported by the DFG (German Research Foundation) postdoctoral fellowship to A. Subkhangulova (DFG project number SU 1131/1-1).

## Competing interests

The authors declare no competing interests.

---

**Figure 1 supplement 1.**
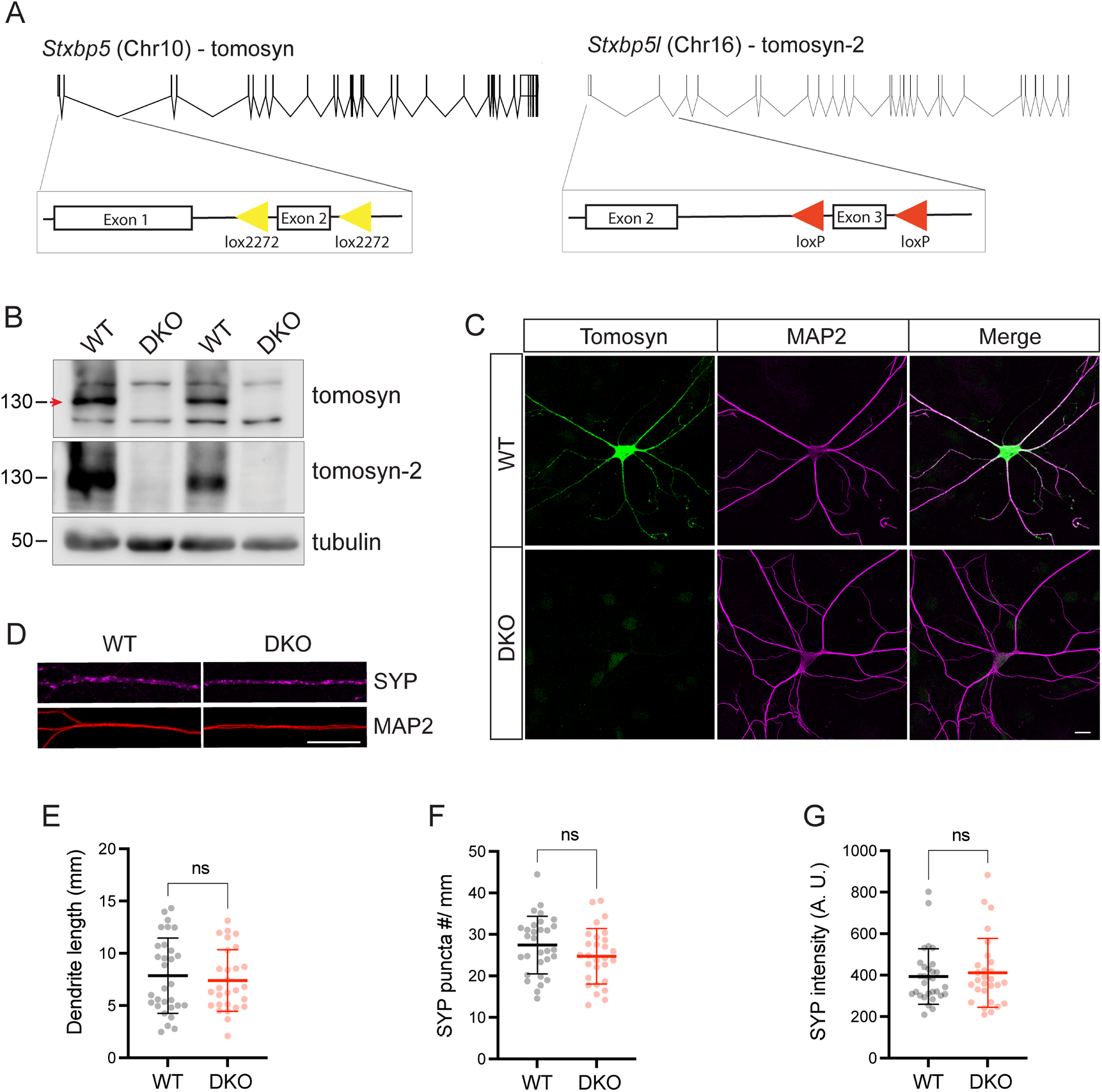
Validation of the mouse model with the conditional deletion of tomosyn and tomosyn-2. (A) Schematic depiction of mouse *Stxbp5* (encoding tomosyn) and *Stxbp5l* (encoding tomosyn-2) genes with exons as vertical lines and introns as bridging gaps. To generate conditional double knockout (DKO) of *Stxbp5/5l*, exon 2 of *Stxbp5* and exon 3 of *Stxbp5l* were flanked with Lox2272 and LoxP sites, respectively (shown as triangles). Expression of Cre-recombinase results in excision of the flanked exons in both genes, predicted to cause frameshift mutations and nonsense-mediated mRNA decay of gene transcripts. Proportional gene graphics were made using Exon-Intron Graphic Maker developed by Nikhil Bhatla. (B) Western blot showing loss of tomosyn and tomosyn-2 expression in Cre-expressing primary hippocampal neurons (‘DKO’). As control (‘WT’), neurons expressing Cre lacking the DNA-binding domain (ΔCre) were used. Arrow indicates tomosyn, which is flanked by two unspecific bands. Numbers on the left indicate approximate molecular weight of the bands in kDa. (C) Immunostaining of tomosyn and MAP2 showing loss of tomosyn expression in DKO neurons. Scale bar 20 µm. (D) Example cropped images of WT and DKO neurons immunostained for the synaptic marker synaptophysin (‘SYP’) showing normal synapse number and distribution in the KO neurons (quantified in E-G). Scale bar 20 µm. (E) Total length of MAP2-positive neurites per single neuron. (F) Number of synaptophysin puncta normalized to length of MAP2-positive neurites. (G) Mean intensity of synaptophysin signal on MAP2-positive neurites. Data in E-G are shown as mean ± SD and were analyzed using a two-tailed unpaired *t*-test. n=28-30 neurons/ genotype. ns: not significant.

**Figure 1 supplement 2.**
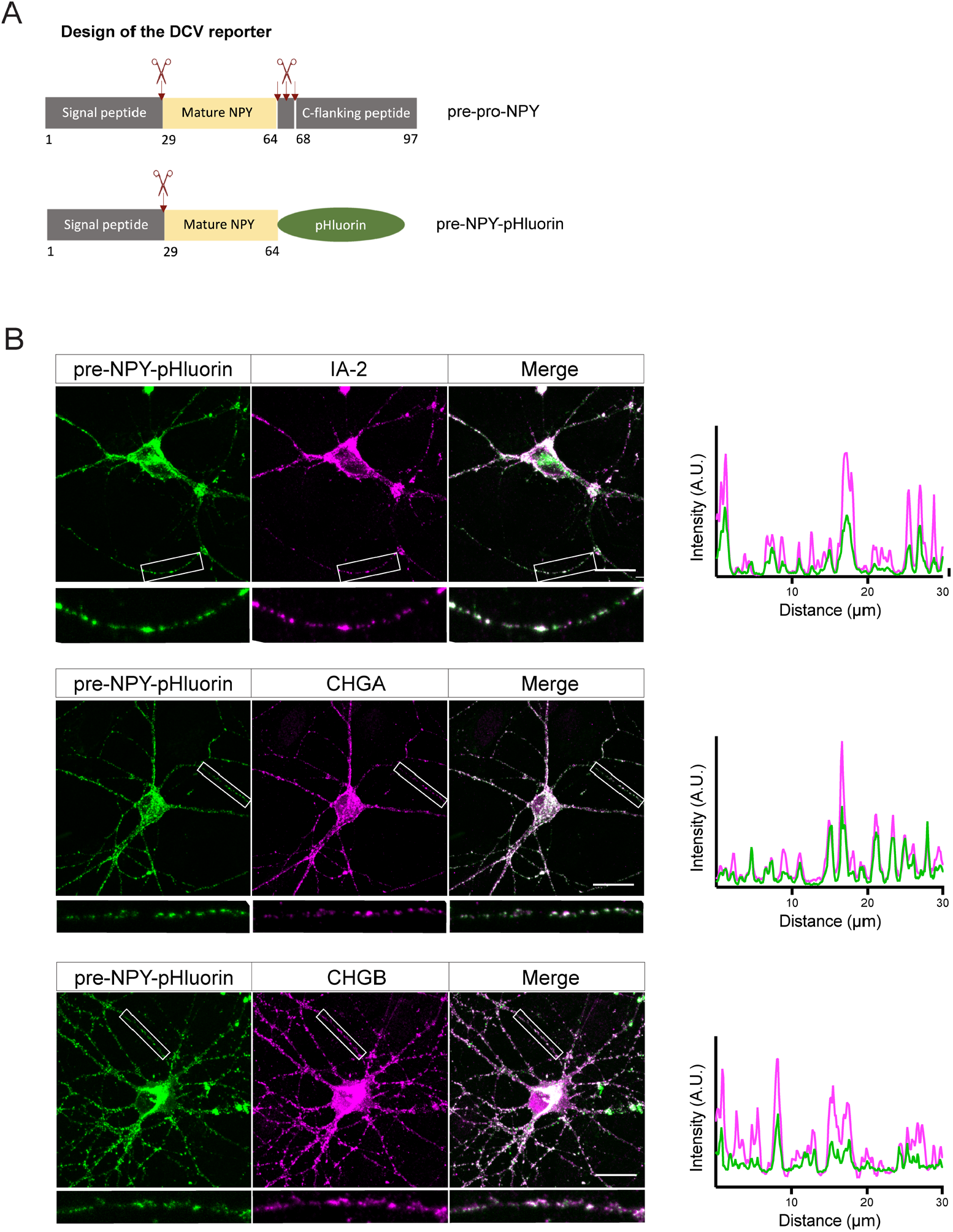
Design and validation of the pHluorin-based DCV fusion reporter. (A) DCV fusion reporter used in this study is based on the human pre-pro-NPY (NPY precursor), which structure is shown in the upper panel with processing sites depicted as scissors. In the DCV fusion reporter, C-flanking peptide (aa 64-97) was replaced by super-ecliptic pHluorin. (B) DCV fusion reporter localizes to endogenous DCVs in cultured neurons, as shown by co-localization of pre-NPY-pHluorin with the endogenous DCV markers IA-2, chromogranin A (CHGA) and chromogranin B (CHGB). White frames indicate zoomed areas used to generate intensity profiles shown on the right. Scale bar 20 µm.

**Figure 1 supplement 3.**
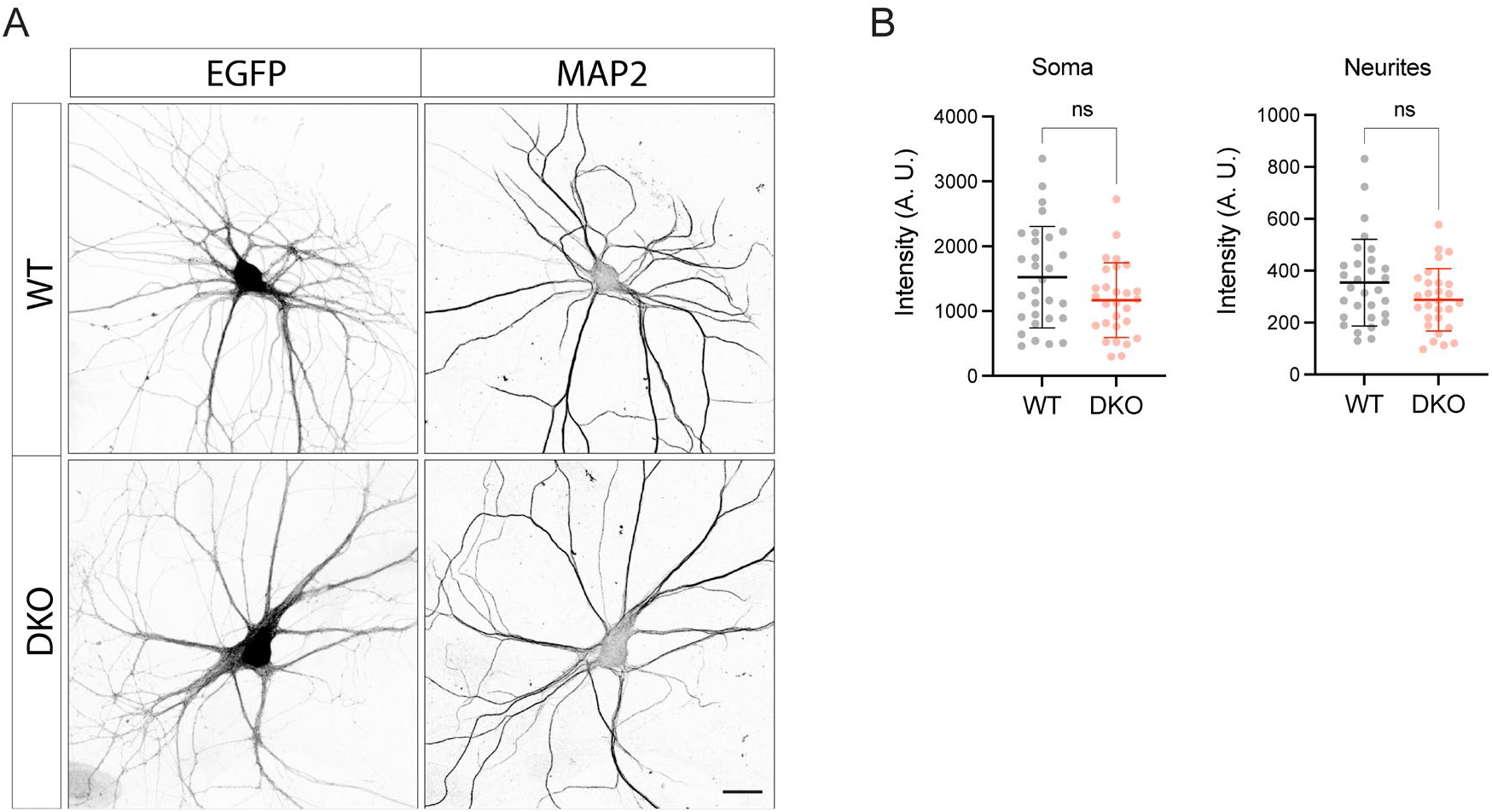
Expression of EGFP under control of synapsin promoter is not affected in DKO neurons. (A) Representative images of EGFP expressed under control of synapsin promoter in fixed WT and DKO neurons. Scale bar 20 µm. (B) Quantification of the mean EGFP intensity in WT and DKO neurons from confocal microscopy images as exemplified in (A). Data are shown as mean ± SD and were analyzed using a two-tailed unpaired *t*-test. n=28-30 neurons/ genotype. ns: not significant.

**Figure 2 supplement 1.**
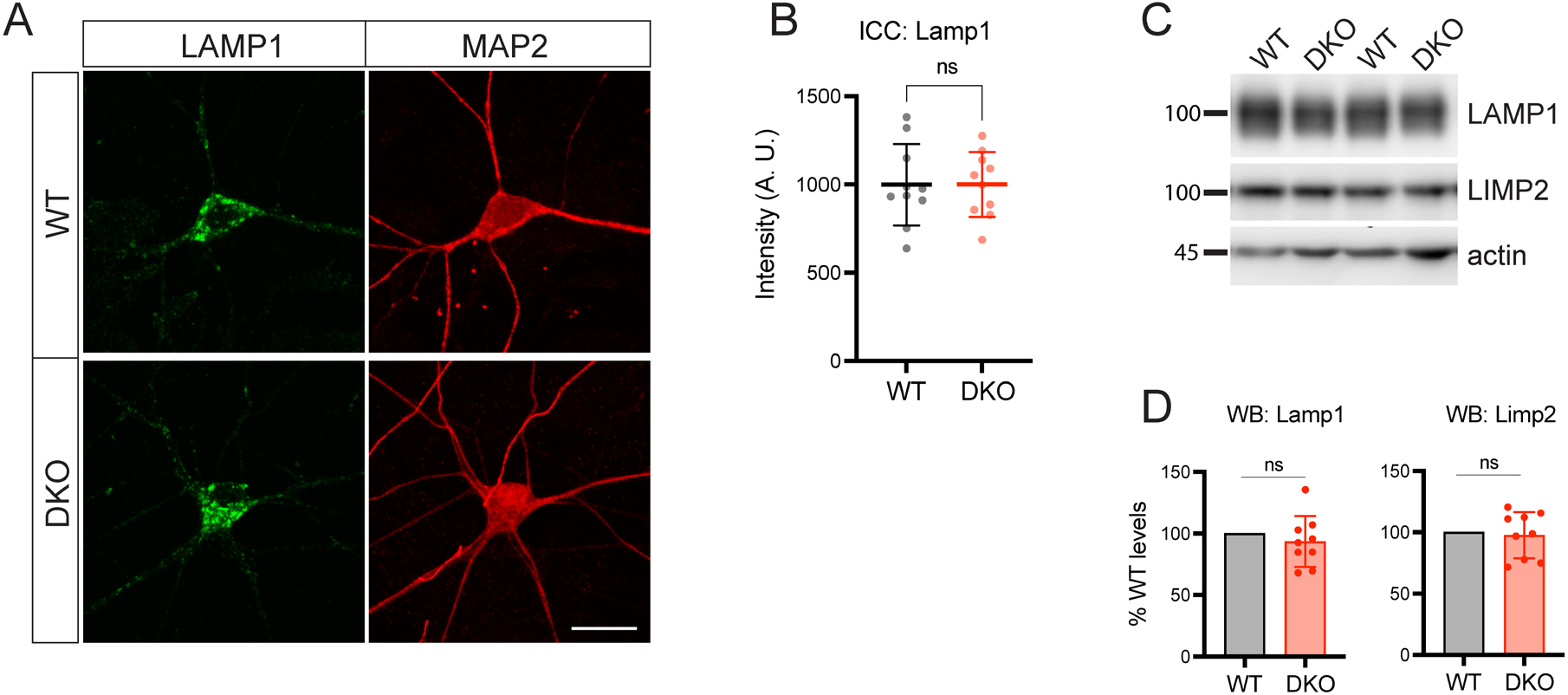
Loss of tomosyns does not affect levels of endo-lysosomal proteins. (A) Representative images of LAMP1 immunostaining in DIV14 neurons. Scale bar 20 µm. (B) Quantification of the mean LAMP1 intensity in WT and DKO neurons from confocal microscopy images as exemplified in (A). Data are shown as mean ± SD and were analyzed using a two-tailed unpaired *t*-test. n=10 neurons/ genotype. ns: not significant. (C) Levels of LAMP1 and LIMP2 as detected by WB are normal in DKO neurons. Equal loading was verified by immunodetection of actin. (D) Quantification of LAMP1 and LIMP2 signal from WB images exemplified in (C). Levels of LAMP1 and LIMP2 in DKO were normalized to WT levels in the corresponding culture. DKO data are presented as mean ± SD and were analyzed using one sample *t*-test. n=9 samples/ genotype from three culture preparations. ns: not significant.

**Figure 5 supplement 1.**
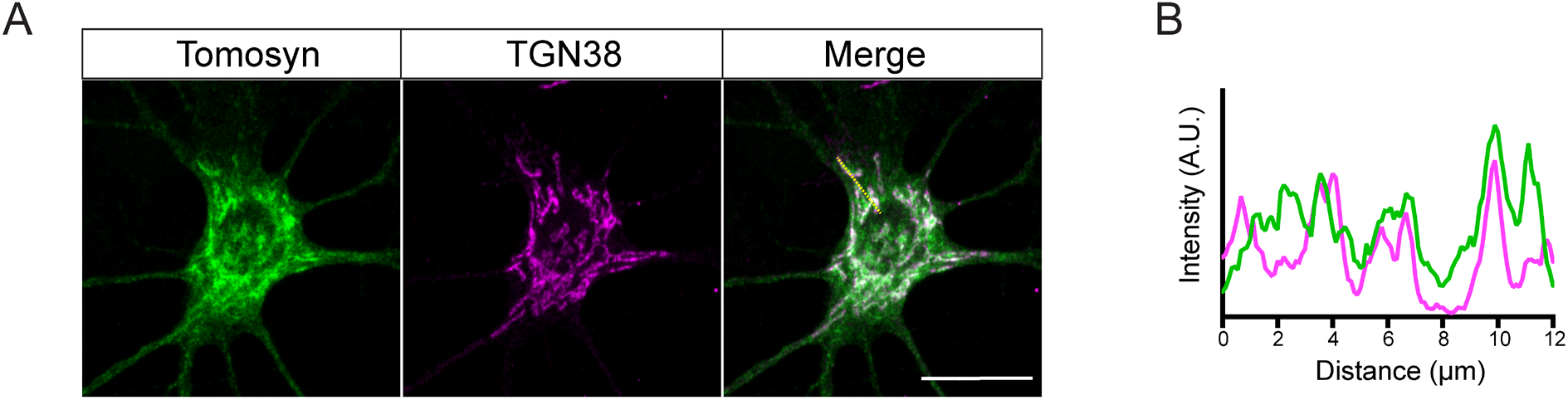
Tomosyn localizes to the TGN. (A) Example confocal microscopy images showing partial co-localization of tomosyn with the TGN marker, TGN38. Scale bar 20 µm. (B) Intensity profile generated from the yellow line indicated in (A) showing partial colocalization of tomosyn and TGN38.

**Figure 5 supplement 2.**
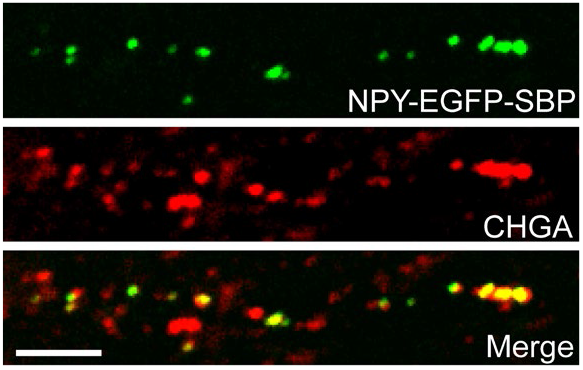
RUSH-generated vesicles contain endogenous DCV marker, CHGA. Example confocal microscopy images showing co-localization of NPY-EGFP-SBP with CHGA. Scale bar 5 µm.

**Figure 6 supplement 1.**
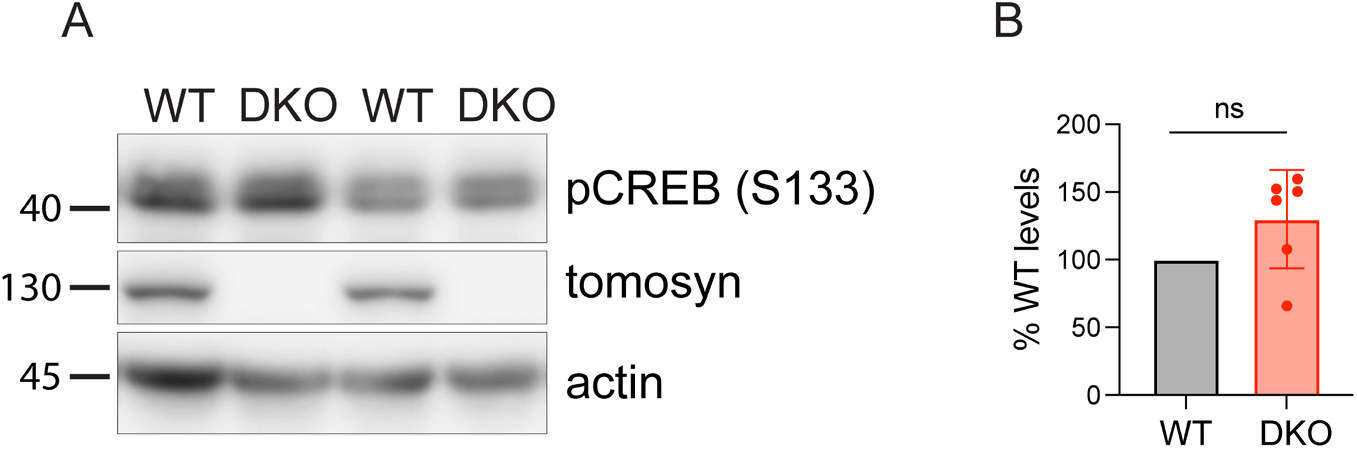
Levels of activated CREB are not affected in DKO neurons. (A) Levels of CREB phosphorylated at S133 as detected by WB are normal in DKO neurons. Equal loading was verified by immunodetection of actin. (B) Quantification of pCREB signal from WB images exemplified in (A). Levels of pCREB in DKO were normalized to WT levels in the corresponding culture. DKO data are presented as mean ± SD and were analyzed using one sample *t*-test. n=6 samples/ genotype from six independent cultures. ns: not significant.

## Notes

### Competing Interest Statement

The authors have declared no competing interest.

### Summary of Updates

This version of the manuscript has figure placement corrected (figures are included in the main text).

